# Host-specific adaptation of *Legionella pneumophila* to single and multiple hosts

**DOI:** 10.1101/2024.11.18.624128

**Authors:** Anaísa B. Moreno, Kiran Paranjape, Martina Cederblom, Elisabeth Kay, Christian Dobre-Lereanu, Dan I. Andersson, Lionel Guy

**Author notes:** These authors contributed equally to the study (Anaísa B. Moreno, Kiran Paranjape).

## Abstract

*Legionella pneumophila* is an endosymbiotic bacterial species able to infect and reproduce in various protist and human hosts. Upon entry in human lungs, they may infect lung macrophages, causing Legionnaires’ disease (LD), an atypical pneumonia, using similar mechanisms as in their protozoan hosts, despite the two hosts being separated by a billion years of evolution. In this study, we used experimental evolution to identify genes conferring host-specificity to *L. pneumophila*. To this end, we passaged *L. pneumophila* in two different hosts - *Acanthamoeba castellanii* and the human macrophage-like cells U937 - separately and by switching between the hosts twice a week for a year. In total, we identified 1518 mutations present in at least 5% of the population at the time of sampling. Half of these were localized in five groups of repeated sequences, likely to be recombination hotspots. Forty-nine mutations were fixed in the 18 populations at the end of the experiment, representing four different groups. The first two groups involve adaptation to the specific selection conditions, including (i) two specific mutations in the 30S ribosomal protein S12 (RpsL) that conferred resistance to streptomycin, to which the bacteria were unexpectedly exposed to during serial passage and (ii) two mutations, one in the 30S ribosomal protein S4 (RpsD), and one in the chaperonin GroES, both of which are likely to be fitness-restoring compensatory mutations to the original RpsL mutations. Two more interesting groups of mutations included (iii) mutations in 4 different strain-specific genes involved in LPS synthesis, found only in the lineages passaged with *A. castellanii* and (iv) mutations in the gene coding for LerC, a key regulator of protein effector expression, which was independently mutated in 6 lineages grown in presence of the U937 cells. We propose that the mutations degrading the function of the regulator LerC improve the fitness of *L. pneumophila* in human-derived cells, and that modifications in the LPS are beneficial for growth in *A. castellanii*. This study is a first step in further investigating determinants of host specificity in *L. pneumophila*.

## Introduction

To date, 65 species of *Legionella* have been discovered, half of which can cause disease in humans. Globally, 85% of cases of legionellosis are caused by *Legionella pneumophila* - with the exception of Australia and New Zealand where 30% of cases are caused by *Legionella longbeachae* (Fields et al. 2002; Den Boer and Yzerman 2004). *Legionella* causes two forms of legionellosis in humans – Legionnaires’ disease (LD) and Pontiac fever. LD is presented as a severe pneumonia, with mortality of up to 15%, whilst Pontiac fever induces a milder flu-like illness. The worldwide occurrence of LD is difficult to calculate due to misdiagnosis and underreporting, but it has been established as a major contributor to both community- and hospital-acquired pneumonia (Falcó et al. 1991; Diederen 2008; Lau and Ashbolt 2009).

*Legionella* has a long history of adaptation to eukaryotes. The order encompassing *Legionella* and the related pathogen *Coxiella*, among others, is believed to have started colonizing eukaryotes almost 2 billions years ago (Hugoson et al. 2022). The ability of *Legionella* to infect humans has been associated with its long history of co-evolution with protozoans. *Legionella* is able to control its host’s cellular processes, such as vesicle trafficking. However, beyond the processes that are shared and highly conserved among all eukaryotes, *Legionella* can also control others that are absent in the protists it frequently encounters (Park et al. 2020). It is unlikely that *Legionella* has evolved these abilities simply from accidental transmission to humans and man-made aquatic environments in the past century, especially because humans are presumably evolutionary dead-ends for *Legionella*, as it doesn’t spread from patient to patient. The human host is not frequently met by *Legionella*, and specific in-patient mutations occur, despite the short infection cycle (Leenheer et al. 2023). This leads to a hypothesis where metazoan species are also natural hosts of *Legionella*, and these have been instrumental in furthering the evolution of *Legionella* to infect human macrophages (Best and Abu Kwaik 2019).

The life cycle of *Legionella* replication is similar in all eukaryotic cells, despite some host-specific differences. It is characterized by two distinct phases, replicative and transmissive (Oliva et al. 2018). Upon entry into the host cell, *Legionella* evades phagosome-lysosome fusion by forming a *Legionella*-containing vacuole (LCV). Shortly after the LCV is formed, it recruits host mitochondria and rough endoplasmic reticulum to surround its surface. In the LCV, *Legionella* undergoes exponential growth - the replicative stage. When the host’s nutrients are exhausted, a new stage is triggered – the transmissive stage. There, *Legionella* is released from the LCV to the host’s cytosol, before exiting the cell using several mechanisms. In this transmissive state *Legionella* is able to infect alveolar macrophages and cause disease in humans. In laboratory conditions, *Legionella* also displays these replicative and transmissive characteristics, during the exponential and stationary phases, respectively (Richards et al. 2013; Oliva et al. 2018).

The capacity of *Legionella* to evade lysosomes and modulate cellular processes in the host is conferred by the Dot/Icm system (Isberg et al. 2009). The latter is classified as a type IVB secretion system (T4BSS), and has two groups of loci - the *dot* (defect in organelle trafficking) and the *icm* (intracellular multiplication). In *Legionella* this secretion system injects over 300 effector proteins into the host cell. These effectors play a role in entry of *Legionella* into the host cell, formation of LCV, and exit from the host (Ensminger 2016). They also manipulate various cellular processes such as vesicle trafficking, protein translation, and ubiquitination pathways. About 10% of the coding capacity of *Legionella pneumophila* is used to express these effector proteins (Richards et al. 2013).

A large number of effector proteins are functionally redundant. This has been illustrated by experiments where deleting 31% of the effectors did not cause significant impairment of intracellular replication of *Legionella* in mouse macrophages (O’Connor et al. 2011). Many effector proteins contain eukaryotic domains which are thought to facilitate modulation of host cells (Hubber and Roy 2010). These domains can combine to make new effectors, which could be one of the reasons why *Legionella* can infect such a wide range of hosts (Gomez-Valero et al. 2019).

An important part of *Legionella*’s infection cycle is the coordination and regulation of effector expression where four two-component systems (TCS) are central (Segal 2014). Together, they regulate the expression of over 100 genes (Feldheim et al. 2018) by sensing environmental stimuli and transducing the signal into the cell, often via an autophosphorylation mechanism. By timing the expression of the right effectors at the right time, they probably play an important role in host specificity.

The broad host-range of *Legionella* is thought to have arisen from its lifestyle of host cycling, which selects against mutations that may decrease its fitness in the many protozoans it naturally infects. This was tested in an experimental evolution set-up, where *Legionella* was passaged in mouse macrophages for a year. The results showed rapid parallel evolution of *Legionella*, and adaptive mutations that resulted in improved replication in macrophages when compared to ancestral strains (Ensminger et al. 2012). Another evolution experiment involving *Legionella* and amoebae is the spontaneous endosymbiosis discovered by Jeon and Lorch (1967). Following an unexpected infection of one of their cultures of *Amoeba proteus* by an unknown bacterium (X-bacterium, now renamed ‘*Ca.* Legionella jeonii’), a few amoebae survived, but kept the bacterium, which turned out to be essential for the amoeba upon subsequent cultivation.

In this study, we aimed at exploring the selective pressures imposed on the multi-host endosymbiont *L. pneumophila*. By using experimental evolution, we sought to identify genetic determinants of host-specificity in this bacterium. We cycled *L. pneumophila* in two different hosts separately and by repeated host switching. After one year of biweekly passaging, we sequenced the evolved lineages at population levels and tested the fitness effects of the most frequent genotypes.

## Results

### Distribution of mutations

Each of the six lineages for all three hosts were passaged in their hosts for up to one year and sequenced at population level at two or three time points (**Supplementary Table 1**). All lineages passaged with human macrophage-derived cells (U937 cells, abbreviated as Mac) were sequenced at all three timepoints, while for the other two conditions, passaged with *A. castellanii* (Acas) and switching between the two hosts (alternation, Alt), only one population was sequenced at one of the two later time points.

We identified in total 1518 mutations that exceeded a 5% frequency in any population, after removing low-frequency mutations (<20%) in one sample (Alt_D, t1) which was contaminated with another strain of *L. pneumophila*. The number of mutations reaching 5% frequency ranged from 4 to 102 per population and time point, with a median of 28.5 (**Supplementary Figure 1**). The number of mutations appeared more variable and on average slightly higher at time point 1 than 3 (**Supplementary Figure 2A**). The fraction of non-fixed mutations is also higher at time point 1 than 3 (**Supplementary Figure 2B**). Small indels accounted for only 4% of the mutations (n = 62). As expected, among SNPs, transitions (987, 68%) were more frequent than transversions (468, 32%). The number of AT −> GC mutations (N = 599, 50.4%) was very similar to the reverse (GC −> AT; N = 589, 49.6%). The numbers were also very similar when excluding repeated regions and recombination hotspots prone to recombination (GC->AT: 270, 50.6%; AT->GC: 264, 49.4%). Very few mutations (15) were found on the *L. pneumophila* Paris pLPP plasmid. All of them appeared with frequencies < 15%. All further analysis focused on the chromosomal mutations.

### Mutational hotspots

Mutations were not uniformly distributed along the chromosome, and many clustered around several hotspots (**Supplementary Figure 3**). In total, mutations in these hotspots accounted for 751 or 49.4% of all mutations, but notably none of them were fixed. Eleven of these hotspots corresponded to intergenic regions upstream or downstream of GIY-YIG nuclease family protein, which is a homing endonuclease/mobile element (Dunin-Horkawicz et al. 2006). In total, 97 mutations occurred in these 11 hotspots. A second hotspot for mutations was a gene corresponding to locus tag lpp1100, coding for an ankyrin repeat-containing protein, which displayed 142 mutations. The ankyrin repeats (19 repeats of 35 residues, **Supplementary Figure S4**) are presumably different enough so that the variants that were called by BRESEQ represent true mutations and not mapping errors. The third hotspot for mutations was one of the five tetratricopeptide repeat (TPR; 19 mutations) proteins present in *L. pneumophila* str. Paris, lpp2912. The fourth hotspot was a repeated protein containing a domain of unknown function (DUF1566; 36 mutations) which is widely present in *Gammaproteobacteria*. Finally, the fifth group of hotspots was constituted of three occurrences of a larger segment containing a transposase and a type II-like restriction endonuclease (DUF559; 457 mutations).

### Frequent and fixed mutations

The frequency of variants varied greatly, but in total 20, 35 and 36 mutations appeared fixed after t1, t2 and t3, respectively; this corresponded to 1.2, 2.7 and 2.8 mutations per population. After the last time point (t3 for Acas and Mac, and t2 for most of the Alt populations), 10, 22, and 15 mutations were fixed in Acas, Mac, and Alt conditions, respectively. In some populations (e.g. Mac_A and Alt_D at t1, Acas_D and Mac_C at t3; **Supplementary Figure 2**), groups of variants were found at a higher frequency but were not fixed, suggesting two sub-populations with distinct mutations co-existing in the same culture.

In addition to the recombination hot-spots regions mentioned above, other genes were also mutated several times with higher frequencies (**Figure 1**). Most fixed mutations (92.3%) were small indels or non-synonymous mutations, while intergenic or synonymous mutations were more common at lower frequencies (**Table 1**), as expected if genetic drift is not prevalent. Remarkably, the number of synonymous mutations dropped from 20.1% among all mutations to only 2.2% (n = 2) among fixed mutations. Many non-synonymous mutations occurred independently in the same gene (e.g. *rpsL*, *lerC*, lpp0832) (**Figure 1**).

**Figure 1:**
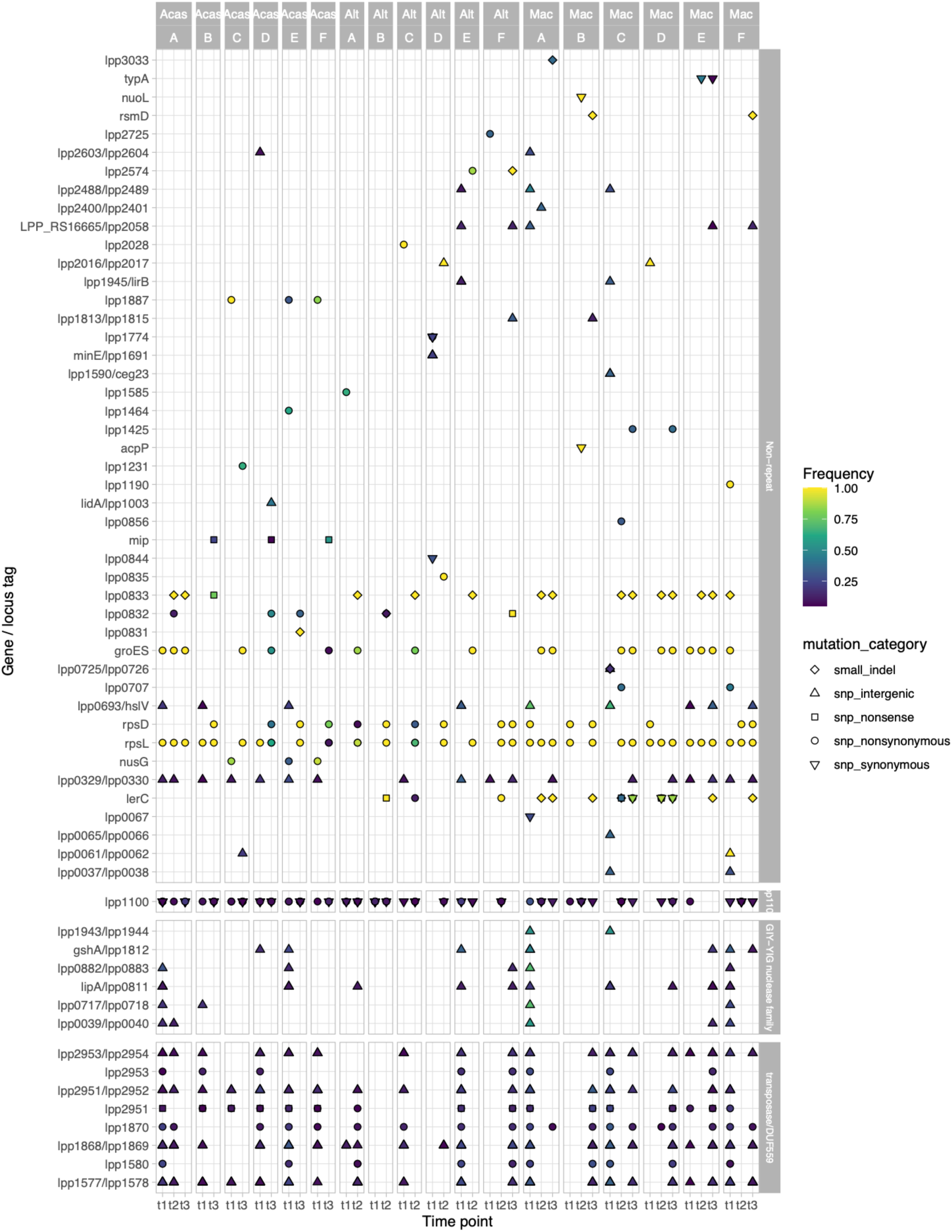
Frequency of mutations appearing at least once within a gene or an intergenic space, with a frequency in the population of at least 25%. Mutations occurring within a gene are labeled with the locus tag or the gene name (if available). Mutations occurring in an intergenic space are labeled with the two bordering gene names or locus tags. If a gene/intergenic space is mutated several times, with different types of mutations (e.g. *lerC*), the different symbols are stapled on top of each other. The shape of the points corresponds to the type of mutation. The color of each point corresponds to the frequency of the mutation. Yellow dots are fixed mutations. Mutations occurring in non-repetitive regions are shown at the top, and repetitive regions at the bottom.

**Table 1:** Number and frequency of mutations per category. Mutations were counted for all three time points. The three pairs of columns represent the number and percentage of mutations with frequency over 5%, with frequency over 25%, and fixed in the population, respectively.

| Mutation category | N (>5%) | % | N (> 25%) | % | N (fixed) | % |
| --- | --- | --- | --- | --- | --- | --- |
| Small indel | 62 | 4.1 | 29 | 3.7 | 23 | 25.3 |
| SNP intergenic | 611 | 40.4 | 404 | 52.1 | 3 | 3.3 |
| SNP nonsense | 42 | 2.8 | 21 | 2.7 | 2 | 2.2 |
| SNP non-synonymous | 492 | 32.6 | 240 | 30.9 | 61 | 67.0 |
| SNP synonymous | 304 | 20.1 | 82 | 10.6 | 2 | 2.2 |
| *Total* | 1511 | 100.0 | 776 | 100.0 | 91 | 100.0 |

In total, 48 mutations (representing 13 distinct mutations in 11 different genes) were fixed in the population after the last time point (**Table 2**). This corresponds to about a year of passaging in the three conditions: ~450 generations in U937 cells (Mac, t3), ~480 generations in the switching condition (Alt, t2), and ~800 generations in *A. castellanii* (Acas, t3) (**Supplementary Table 1**).

**Table 2:**
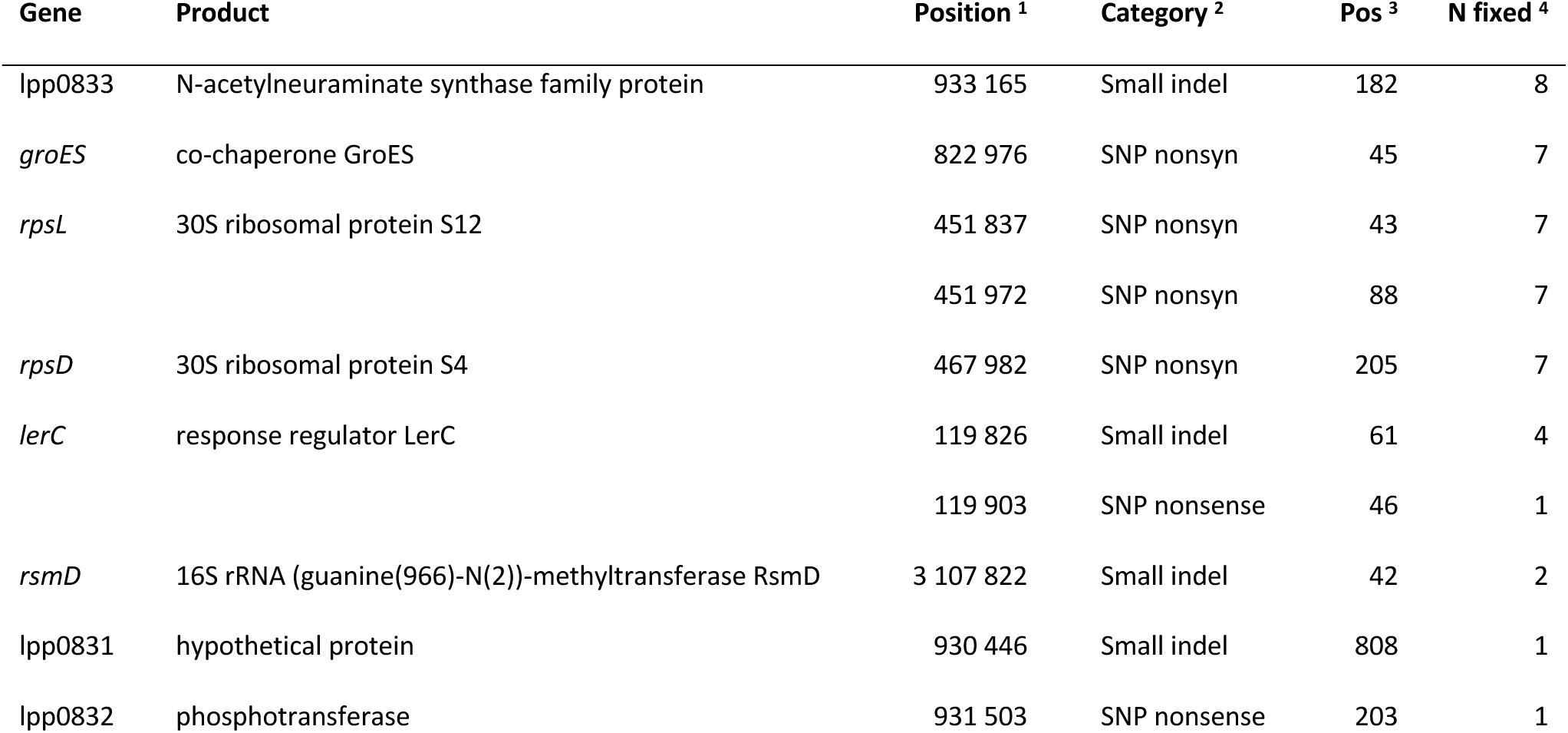
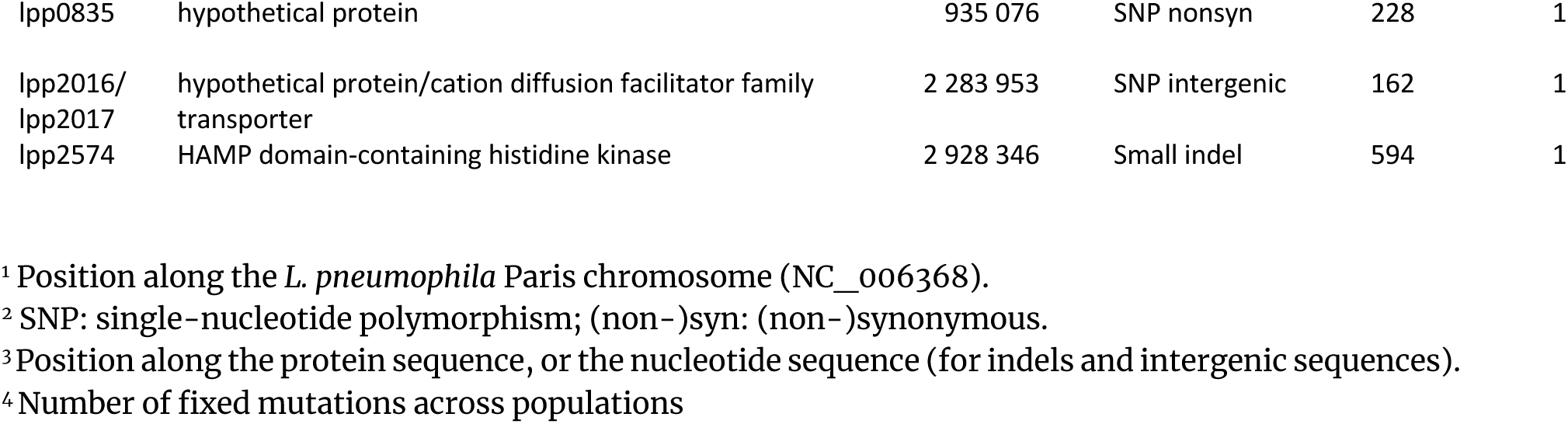
Fixed mutations. Mutations shown are those fixed at the last time point for each lineage.

As expected, most fixed mutations were non-synonymous or nonsense mutations, or small intragenic indels (**Figure 2**). Two synonymous mutations (in *acpP* and *nuolL*, **Figure 1**) appeared to be fixed in the Mac_B population but were absent at the third time point. One intergenic mutation was found, 162 respectively 187 nucleotides upstream of two genes, lpp2016 and lpp2017 (**Figure 2**). The latter gene is, according to in-silico predictions (Price et al. 2005), the first in a two-gene operon (lpp2017 and lpp2018), both involved in ion transport. The former gene, lpp2016, has no described function, and appears to be strain-specific in *L. pneumophila*: it is present in subspecies *pascullei* and in strains Paris, Alcoy and Corby but absent in Philadelphia (Altschul et al. 1997).

**Figure 2:**
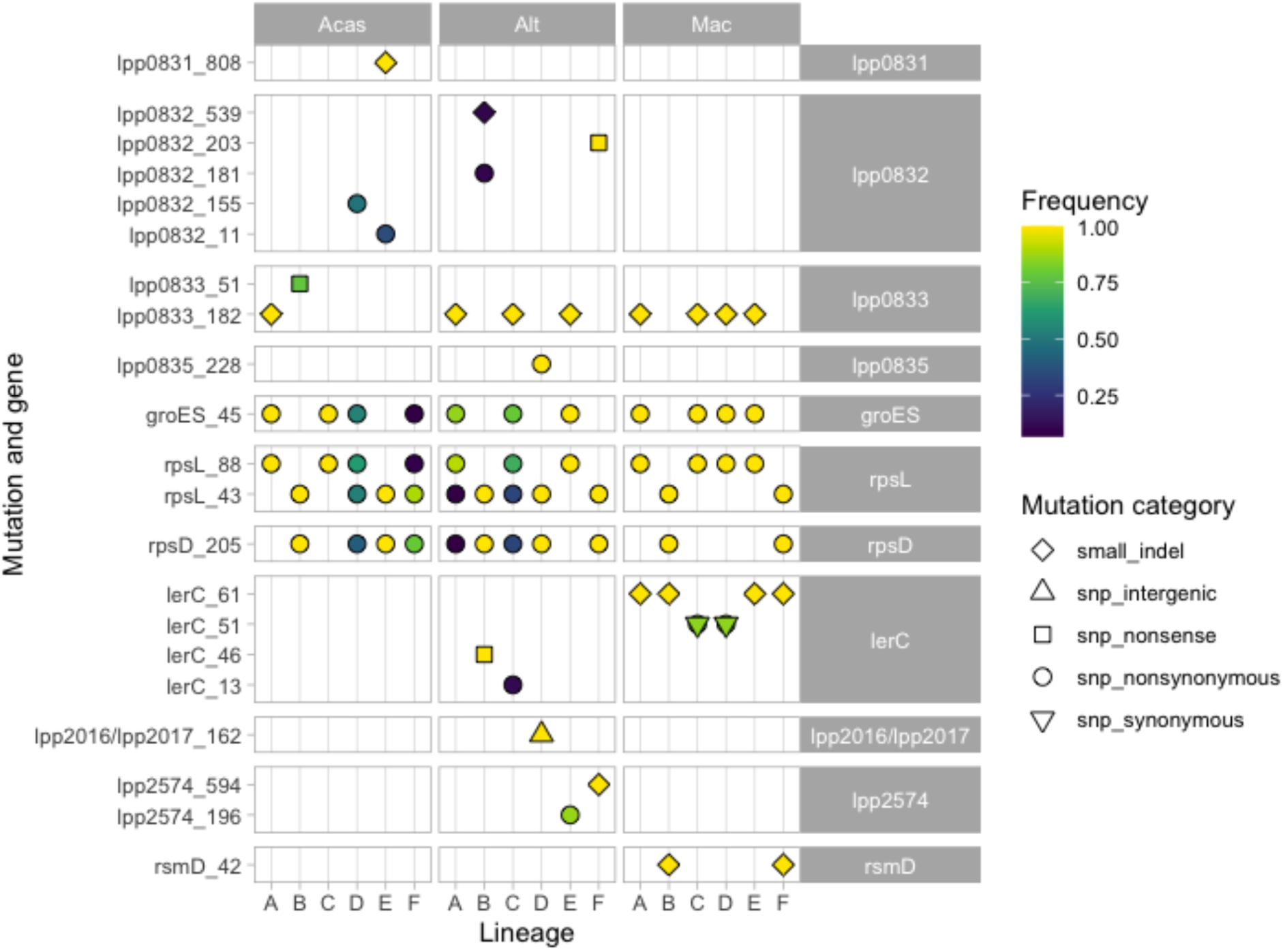
Genes with fixed mutations at the last sequenced time point under the three conditions. Mutations occurring within a gene are labeled with the locus tag or the gene name (if available) on the right side of the figure; mutations occurring in the same gene at different positions (*rpsL*, *lerC*) are shown on different lines, and the position of the mutation is shown on the left side of the panel. For in-gene SNP mutations, the position of the amino-acid whose codon has been mutated is indicated. For the intergenic mutation, the position is the number of nucleotides after the upstream gene; for the small indels, the position refers to the nucleotide position within the gene for the start of the indel. Mutations occurring in an intergenic space are labeled with the two bordering gene names or locus tags. The shape of the points corresponds to the type of mutation. The lerC_51 mutation is a double mutation in the same codon, at the second and third codon position, resulting in a non-synonymous mutation. The color of each point corresponds to the frequency of the mutation. Yellow points are fixed mutations.

### Two pairs of mutations in ribosomal protein and chaperonin genes are mutually exclusive

The distribution of mutations showed two distinct patterns of mutations, both of which were present under all three conditions (**Figure 2**). In one (referred to as RpsL43/RpsD), the 30S ribosomal protein S12 RpsL was mutated at position 43, where the lysine is replaced by a threonine (K43T), while the 30S ribosomal protein S4 RpsD is mutated at position 205, where a serine replaces a tyrosine (S205Y). In the other pattern (RpsL88/GroES), RpsL has another mutation, where the lysine at position 88 is replaced by an arginine (K88R), while the co-chaperone GroES has a mutation at position 45 (A45T). Every population harbored one or the other pair of mutations, or a mix of both, at the last time point. Strikingly, these two pairs of mutations were never seen together on a single isolate. In the four cases where both patterns were present (Acas_D, Acas_F, Alt_A, and Alt_C; **Figure 2**), the frequencies of the two mutations within the pattern were very similar, and the sum of the two patterns roughly equals one. An inspection of the reads spanning both positions in the *rpsL* gene revealed that no single read displayed both mutations, but that all reads carried one.

The two mutations in RpsL (K43T and K88R) have previously been shown to provide resistance to streptomycin in *L. pneumophila* (Rao et al. 2013). We determined the minimal inhibitory concentration to be over 1024 μg/ml in both cases. For both mutations, a halo zone was observed on plates (**Supplementary Figure S5**) but the halo was larger in the K88R-carrying mutant (down to ~96 μg/ml) than in the K43T-carrying mutation (down to ~768 μg/ml).

### Mutations in the LPS synthesis gene cluster

Four linked genes were harboring fixed mutations: lpp0831 (a small indel), lpp0832 (one fixed nonsense mutation and four non-fixed mutations: one small indel and three non-synonymous mutations), lpp0833 (one fixed small indel in many populations, as well as a non-fixed nonsense mutation in Acas_B), and lpp0835 (one non-synonymous mutation) (**Figure 2**). These four genes belong to the variable segment of the lipopolysaccharide (LPS) synthesis gene cluster. The mutation in lpp0833 was fixed in 8 out of 18 lineages, and only found in populations which harbored the RpsL88/GroES genotype. In two of these lineages (Alt_A and Alt_D), though, the lpp0833 mutation was found in 100% of the population whereas the two genotypes RpsL88/GroES and Rpsl43/RpsD coexisted. Thus, the lpp0833 was found also in the Rpsl43/RpsD background in these two lineages. The other mutations in the LPS synthesis operon were found exclusively in the lineages passaged in Acas or during switching conditions.

### Mutations in LerC, a response regulator of the effector regulatory network

Four distinct mutations in LerC were found at the last time point: a A13D non-synonymous change (lerC_13), a 13-nt insertion after nucleotide position 61 in the gene (codon position 21; lerC_61), a C46* nonsense mutation (lerC_46), and a double SNP, resulting in a non-synonymous V51G mutation (lerC_51). All Mac lineages harbored one LerC mutation, and it was fixed in four of them; two of the Alt lineages harbored one (B and C, fixed in B), but none was found in the Acas lineages (**Figure 1, 2**). All these mutations were absent at the first time point.

The two non-synonymous lerC_13 and lerC_51 mutations seemed to have little effect on the structure of the LerC protein (**Supplementary Figure 6A and D**). The lerC_46 and lerC_61 mutations yielded a short peptide at the N-terminal (45 and 20 residues, respectively). However, longer proteins (88 and 122 residues, respectively) can presumably be translated from alternative start codons, to the end of the protein. The lerC_61 mutant protein lacks the first two β-strands and the first α-helix (**Supplementary Figure S6B**), while the lerC_46 mutant protein lacks an additional α-helix and the conserved aspartic acid residue at position 53 (**Supplementary Figure S6D**), which is essential for the phosphorylation and function of LerC (Feldheim et al. 2018).

### Other mutations

Three other regions harbored fixed mutations at the last time point (**Table 2**; **Figure 2**). The first one was an intergenic region, upstream of both lpp2016 and lpp2017, in a location where the mutation could play a role in the regulation of either or both genes. The second was the gene lpp2574, which encodes a HAMP domain-containing histidine kinase. Two different mutations were found in this gene, a 1-nt insertion in the middle of the gene (lpp2574_594), fixed in Alt_F, and a non-synonymous R196S (lpp2574_196), present in 86% of the Alt_E population. The third one was the gene coding for the rRNA methyltransferase RsmD (**Figure 3**), which methylates the guanine 66 of the 16S ribosomal RNA during ribosome assembly. The effect of this mutation (rsmD_42) is unclear but is likely to result in a loss-of-function mutation, as the mutation consists in a 1-nt deletion early in the gene.

**Figure 3:**
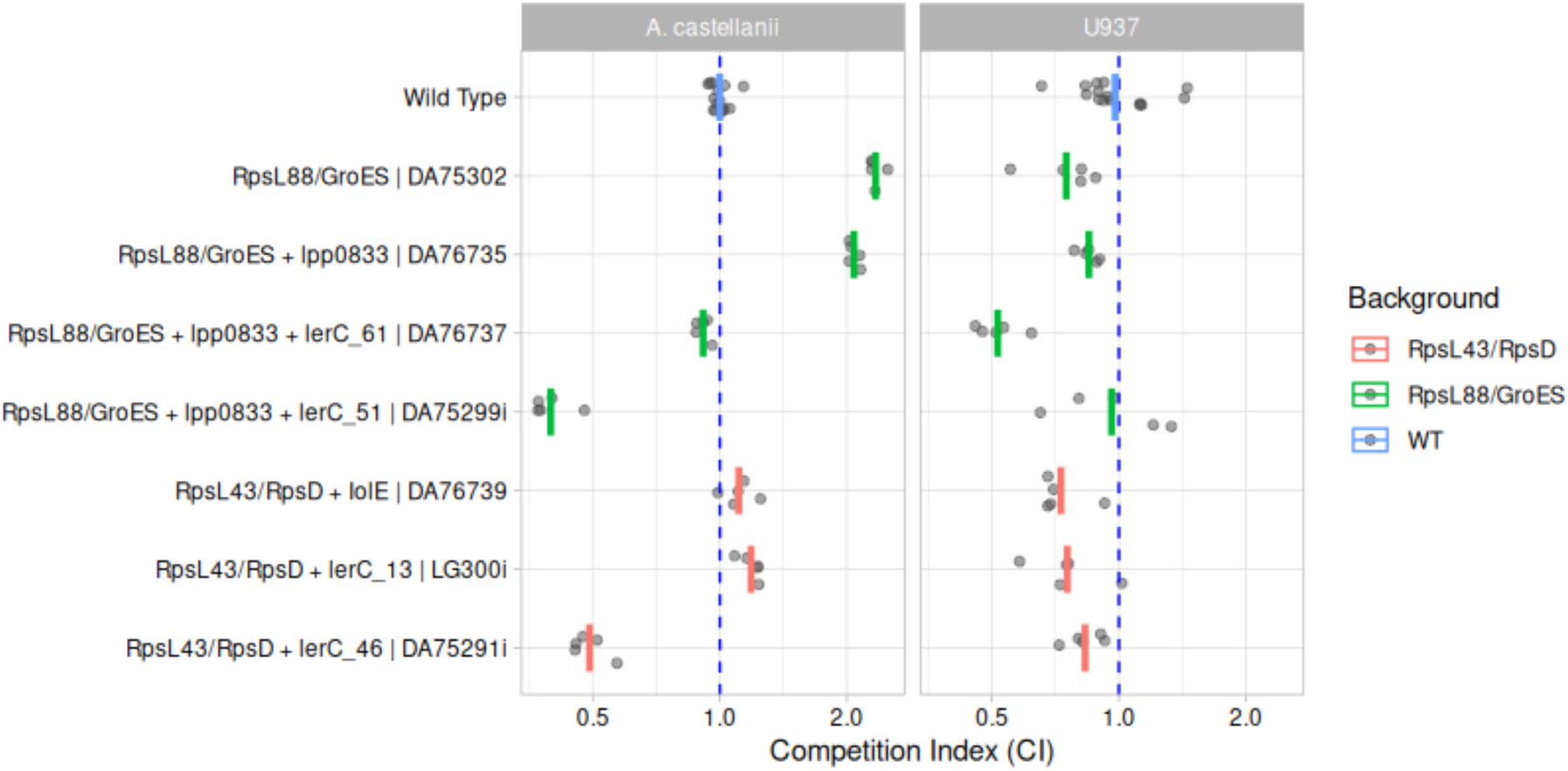
Competition index (CI) of different isolated strains evolved during the evolution experiment. Evolved strains of *L. pneumophila* Paris were competed in the U937-like macrophage cells and the amoeba *A. castellanii* against the ancestral strain. Each replicate is represented by a grey dot and the average is shown as a red bar. The blue dashed line is set at CI=1 (evolved clones grow as much as the ancestors).

### Fitness effects

To estimate the fitness effects of the mutations occurring in the experiment, clones were isolated from populations and whole-genome sequenced. Their growth was compared to their ancestors’ growth, in both host cell types (U937 cells or *A. castellanii*). Since the two fluorescent markers (SYFP2 and dTomato) differ in many aspects (intrinsic effect on organism fitness, brightness, folding time, half-life, etc.), it is not possible to infer fitness differences between two clones carrying different fluorescent markers directly from competition experiments. To correct for this, a parallel experiment was run, competing the two ancestors, isogenic except for the fluorescent marker. The fitness of the evolved clone, competed against the ancestor with the other marker, can be compared to the fitness of the ancestor with the same marker, competed against the ancestor with the other marker. Two methods to estimate fitness were evaluated: the first one assessed growth rates by fitting splines to the growth curves, the second one estimated total growth by comparing the fluorescence in stationary phase to the fluorescence early in the experiment. The first method proved difficult to reproduce consistently, and the second method, which calculates a Competition Index (CI), similar to the method used by Ensminger et al. (2012), was preferred.

CI were calculated between the beginning of incubation (0h) and 64h of infection. This timeframe represented the beginning and end of infections in our experimental setup (see **Supplementary Figure S7**). Since a MOI of 10 was used, representing 10 bacterial CFUs for 1 host cell, it allowed for only one replication cycle. A CI of 1 is equivalent to a neutral effect in terms of fitness between the evolved and the ancestral strain. CI measures the total production of cells during one cycle of replication of *Legionella* within its host. It captures the growth rate only indirectly, and it does not capture differences in how bacteria exit the host cell. The results of competition experiments are shown in **Figure 3**.

The gene *rpsL*, encoding the 30S ribosomal protein S12, was mutated in all populations, conferring streptomycin resistance to *L. pneumophila*. Two distinct mutations (at position 43 and 88) were identified, and each was accompanied by another mutation. The RpsL43/RpsD double mutation is found in 11 out of 18 lineages, and is fixed in 7 out of them. The other double mutation conferring streptomycin resistance, RpsL88/GroES, is equally widespread, found in 11 out 18 lineages, and fixed in 7. One isolate with the second genotype (Rpsl88/GroES; DA75302) showed a strong advantage in competitions in amoebae, but a slight disadvantage in macrophages (CIs of 2.34 and 0.76, respectively) (**Figure 3A**). None of our isolates had only the RpsL43/RpsD genotype, but one (DA76739) had these two mutations and an extra G241V non-synonymous mutation in the IolE protein, involved in myo-inositol metabolism. This isolate showed a similar trend, but not as pronounced, as the RpsL88/GroES mutant: it grew slightly better than the wild type in amoebae but worse in macrophages (CI values of 1.11 and 0.73, respectively) (**Figure 3B**).

Another gene with several independent mutations was *lerC*, which encodes a single-domain response regulator, involved in modulating the expression of the protein effectors translocated into the host cytoplasm (Feldheim et al. 2018). Four isolates contained mutations in their *lerC* gene: DA75291i with a nonsense mutation (lerC_46, C46*); DA75299i with a non-synonymous double mutation (lerC_51, V51G); LG300i with another non-synonymous mutation (lerC_13, A13D), and DA76737 with a 13-bp deletion (lerC_61). The results of the competition with the ancestors are shown in **Figure 3B**, while the predicted structure of the mutants is shown in **Supplementary Figure S6**. All these isolates also contained streptomycin resistance-conferring mutations: DA75291i (lerC_46, C46*) and LG300i (lerC_13, A13D) also harbor the double mutation RpsL43/RpsD, while DA75299i (lerC_51, V51G) and DA76737 (lerC_61, 13-bp deletion) harbor the other double mutation RpsL88/GroES. The latter two isolates, DA75299i and DA76737, also harbor an extra mutation in lpp0833, involved in LPS synthesis. The effects of the mutations are thus compounded in the fitness experiments.

In *A. castellanii* cells, the RpsL88/GroES mutations provided the largest fitness gain (DA75302; CI = 2.34). Adding a mutation in the lpp0833 (involved in LPS synthesis), the fitness was larger than for the wild type, but slightly lower than the latter mutant (DA75735; CI = 2.08). The further addition of the 13-bp deletion in LerC (lerC_61) results in a fitness similar to the wild type (DA75737; CI = 0.91) while the addition of the non-synonymous lerC_51 mutation resulted in a very large fitness drop (DA75299i; CI = 0.40), suggesting that the net effect of both LerC mutations is a large fitness loss. On the other hand, the RpsL43/RpsD (accompanied by the IolE mutation) yielded a very slight fitness gain (DA76739; CI = 1.11). Removing the IolE mutation but adding the non-synonymous lerC_13 mutation resulted in a very similar fitness (LG300i; CI = 1.19), while the addition of the nonsense lerC_46 mutation resulted in a large fitness drop (DA75291i; CI = 0.49) (**Figure 3**). Unexpectedly, although mutations in the LPS synthesis cluster were widespread in populations exposed to *A. castellanii*, the largest fitness increase seem to be provided by the RpsL88/GroES mutations.

In macrophages, the mutations had in general a weaker effect, and most tested mutations resulted in a slight fitness drop. The CI values were on average between 0.75 and 1 (with one exception), and showed a greater variance than in amoebae, making comparisons less reliable. One isolate, DA76737, which has the four mutations (in RpsL88/GroES, in lpp0833, and in lerC_61) that are fixed in two lineages (Mac_A and Mac_E) had a markedly lower fitness compared to the ancestral type, with an average CI of 0.52. This value can be compared to the CI of DA76735, which has the same mutations but lacks the non-synonymous lerC_61 mutation (CI = 0.85). Despite the ubiquity of mutated LerC sequences in populations exposed to macrophages, no LerC mutants seemed to show an increased fitness in macrophages, at least in the experimental setting used here.

### Mutation rates and diversity of streptomycin resistance-conferring mutations

To investigate the likelihood that the same mutations (in particular the ones conferring resistance to streptomycin) occur many times in parallel, we measured mutation rates to streptomycin using fluctuation assays for the three ancestral strains (*L. pneumophila* Paris, and both SYFP2- and dTomato-marked descendants). The results indicated that the mutation rates were low for all three strains (see **Supplementary Table 2**), ranging from 5.5 × 10^-10^ to 7.6 × 10^-10^. Considering an average mutation rate of 6.5 × 10^-10^ and that the bacterial population size is 5 **×** 10^5^ and 5 **×** 10^6^ for the experiments with *A. castellanii* and U937 cells, respectively, there is a 3.3 **×** 10^-4^ and 3.3 **×** 10^-3^ probability that a mutation conferring resistance to streptomycin will appear in one cell at each generation. At t1, which corresponds to 128 and 113 generations (**Supplementary Table 1**), the probability that streptomycin resistance would have occurred at least once in any given Acas and Mac population reaches 4% and 30%, respectively. At t3 (798 and 444 generations) the probabilities were 23% and 76%, respectively. However, the streptomycin resistance was present in half the Acas and 5 out of 6 Mac populations at t1, and in all populations at t3, exceeding the calculated likelihoods, indicating that streptomycin resistance might have introduced by contamination.

To investigate whether the only two RpsL mutations we obtained in our evolution experiment (K43T and K88R) were representative of all streptomycin-resistance conferring mutations, we sequenced the *rpsL* gene of 40 streptomycin-resistant isolates obtained from the fluctuation assays. Although six other mutations were identified, K43T (11 mutations) and K88R (21 mutations) accounted together for 80% of the mutations found in these isolates (n = 40) (**Supplementary Figure S8**).

## Discussion

In this study, we experimentally evolved *Legionella pneumophila* in presence of one of its natural hosts, the amoeba *Acanthamoeba castellanii*, and of the human macrophage-derived cell line U937, to identify host-specific genetic adaptations. We passaged *L. pneumophila* separately in each host and switching between them, twice a week for a year. In the populations exposed to the amoebal host, we found that the LPS-synthesis cluster was probably under selection whereas in the populations exposed to the human macrophage-derived cell lines, LerC, a protein playing a major role in regulating the temporal expression of protein effectors, harbored potential adaptive mutations. We also found that most mutations occur in mutational hotspots, presumably as a result of the high recombination rates in *L. pneumophila*.

### Host-specific adaptations in the LPS-synthesis cluster and in the LerC response regulator

Most *L. pneumophila* populations exposed to *A. castellanii* displayed mutations in genes of the LPS synthesis cluster. The *Legionella* LPS synthesis cluster contains ca. 30 genes (lpg0779-lpg0746 in the Philadelphia strain; lpp0812-lpp843 in Paris) (Thürmer et al.). The genes that were mutated in our evolution experiment (lpp0831, lpp0832, lpp0833 and lpp0835) correspond to ORF 11, 10, 9 and 7, respectively, and belong to the serogroup 1 (SG1)-specific region of the LPS gene cluster (Mérault et al. 2011; Petzold et al. 2013). Whereas lpp0831, lpp0832 and lpp0833 have homologs in other serogroups (albeit with low similarity), lpp0835 is found only in SG1, and has been proposed as a marker for that group (Mérault et al. 2011). The function of the heavily mutated lpp0833 (lpg0768) is not fully understood, but it is homologous to a sialic acid synthetase function (neuAc, neuB family), and is presumably involved in late modifications of the O-antigen (Petzold et al. 2013). The 1-nt deletion in lpp0833 (lpp0833_182) results in an out-of-frame mutation after 182 nucleotides (of >1000), resulting in a very truncated protein. Although functional studies would be required to firmly establish the effect of the truncation, it is likely that it results in a non-functional protein.

The fact that 8 distinct mutations were found in the LPS operon, affecting only the populations in contact with amoebae, strongly suggest that these are adaptive, and beneficial for *Legionella* when infecting the *A. castellanii* strain used in this experiment. One isolate with the 1-nt deletion (lpp0833_182) indeed showed a very strong advantage in competitions inside *A. castellanii*. However, this isolate also carries the RpsL88/GroES genotype, which itself provides an important advantage. It is thus difficult to separate the effect of the two genotypes. Further research in the function of these genes in different hosts might uncover the relationships between LPS and host-specificity, but a reasonable hypothesis is that the LPS obtained in the evolved *L. pneumophila* clones is different to the one in the wild-type. The modified LPS thus potentially lowers the immune response of *A. castellanii*, providing an advantage for this particular *Legionella* strain.

The response regulator LerC is another gene carrying multiple distinct mutations, (namely four), two of which are fixed in 5 populations after the last time point. LerC is involved in the network of regulation of effector proteins (Feldheim et al. 2018). Primary regulation of the expression happens mainly through four distinct two-component systems (TCS), which together regulate over 100 effectors (Segal 2014). Among these, PmrAB regulates directly over 40 effectors (Al-Khodor et al. 2009), while LetAS regulates another set of effectors, via the small RNAs RsmY and RsmZ (Sahr et al. 2009). LerC was found to be a connector between these two systems: it is activated by PmrAB, and inhibits LetAS (Feldheim et al. 2018). The repression of LetAS seems to be growth-phase dependent, as PmrAB is activated under the exponential growth phase, favoring cell divisions inside the *Legionella*-containing vacuole (LCV). At some point, late in the exponential phase, the activation of LerC by PrmAB decreases, which in turn relieves the repression on LetAS, which triggers the cells to enter stationary phase (Feldheim et al. 2018).

In the study describing the function of LerC (Feldheim et al. 2018), the authors noticed that knock-out mutants of *lerC* resulted in various colony types where one of the types took over after a few passages. Sequencing the new dominant type revealed, in addition to the *lerC* mutation, mutations in either *letA* or *letS* that resulted in non-functional versions of the proteins. Although we found several non-functional versions of LerC in our experiment, *letA* and *letS* were not mutated in any of our strains. However, *lerC* mutants were grown on solid medium by Feldheim et al., while our lineages were grown in presence of their hosts only. This indicates that repression of LetAS might be important for growth on solid medium. It should also be noted that Feldheim et al. used *L. pneumophila* JR32, derived from Philadelphia-1, which are separated by ca. 70 000 SNPs from the Paris strain used here (David et al. 2016; Wee et al. 2021): the regulation of effector secretion might be different between the two strains, and could explain why we were able to obtain LerC mutants without the compensating *letAS* mutations.

Mutations in LerC appeared to be generally deleterious in *A. castellanii*, as evidenced by the absence of any LerC mutant in the populations exposed to this host only. This is further supported by competition experiments: compared to the strain with the same background except LerC, the LerC mutants had a lower CI. One strain with the wild-type LerC (DA76735) had a CI of 2, while two other strains, harboring two different LerC mutations (DA76737 with the lerC_61 deletion; DA75299i with the non-synonymous V51G mutation) had much lower CIs, under 1 and under 0.4, respectively. The comparisons for the other two mutants are more complicated to interpret, as there were no isolates with exactly the same background, but the mutant with the lerC_46 nonsense mutation (DA75291i) had a CI of 0.49, more than half than the counterpart (DA76739) which had the wild-type LerC (CI = 1.11) but also another mutation in IolE. The last lerC mutant (LG300i), carrying lerC_13, a nonsynonymous mutation (A13D), had actually a slightly higher CI (1.19) than DA76739 (1.11). In summary, both the distribution, diversity and competition results show that mutations affecting LerC are deleterious for the survival and reproduction of *L. pneumophila* in *A. castellanii*.

On the other hand, the mutations in LerC are likely to be adaptive in human macrophage-derived cells: all six populations passaged in U937 cells only acquired a mutation in the *lerC* gene, which got fixed in four of them. Two populations passaged in alternation in U937 and *A. castellanii* cells showed mutations in LerC, of which one got fixed. The expected advantage in macrophages was only partially confirmed by our competition experiments. The comparison with the wild-type strain is complicated by the fact that the LerC mutants in our collection of isolates all have additional mutations. The most widespread mutation (lerC_61) had a lower CI (0.52) than its counterpart without (0.82), which is quite unexpected. Conversely, the isolate with the lerC_51 mutation (DA75299i), present in the majority of both the Mac_C and Mac_D populations, had a higher CI than its counterpart without (DA7535). The fitness for the other two LerC isolates (lerC_46, and lerC_13; also harboring the RpsL43/RpsD mutations) is again difficult to evaluate, as there are no isolates with only the RpsL43/RpsD mutations. However, compared to the RpsL43/RpsD isolate that also has an IolE mutation, the lerC_46 and lerC_13 isolates have a slightly higher CI. In summary, the pervasive presence of LerC mutants in the populations passaged in human macrophage-derived cells strongly suggests that the wild-type LerC protein is deleterious to *L. pneumophila* Paris when infecting these cells.

In summary, although the results of the competition experiments are somewhat inconclusive, the pervasive presence of LerC mutations in populations exposed to macrophages-like cells but not in populations exposed to *A. castellanii* only, suggests that most LerC mutations are deleterious for growth in *A. castellanii*, while they are likely advantageous for growth in human macrophage-like cells. The variety of mutations found in the lineages exposed to macrophage-like cells, including two that significantly shorten the protein, suggests that a loss or reduction of the LerC function would be beneficial in the human host. The fact that the LerC mutants are also found in lineages where *Legionella* was alternated between the two hosts suggests that the positive selective value in macrophage-like cells is large enough that it overcomes the negative selective values in *A. castellanii*. Thus, LerC is a good candidate for a gene that confers a host-specific advantage, in this case in human cells.

### Adaptation to experimental growth conditions

In addition to the potentially host-specific adaptations, we identified groups of mutations involved in adaptation to the specific growth conditions of our experiment. Among them are two mutations in the ribosomal protein S12 RpsL (K43T and K88R), known to provide resistance to streptomycin (Funatsu and Wittmann 1972). Spontaneous resistance to streptomycin resistance in *Legionella* is a common phenomenon, e.g. in *L. dumoffii* (Hubber et al. 2017) or in many laboratory descendants of the *L. pneumophila* Philadelphia (Berger and Isberg 1993; Rao et al. 2013). Single-step resistance to streptomycin through mutation of *rpsL* has been documented, among others, in *Escherichia coli* (Funatsu and Wittmann 1972) and in *Mycobacterium tuberculosis* (Nair et al. 1993). In both organisms, mutation of codon 43 (42 in *E. coli*) provides resistance to streptomycin, while mutations in codon 88 (87) is found mostly in *E. coli* (Timms et al. 1992). Both mutations have been identified in spontaneous streptomycin mutants in *L. pneumophila* (Rao et al. 2013). Our *Legionella* lineages were never directly exposed to streptomycin, but early in the experiment, both host cell lines were grown in the presence of penicillin-streptomycin, to avoid contaminations. The host cells were thoroughly washed before being infected with the *Legionella* lineages, precisely to prevent exposing *Legionella* to the antibiotic. A possible explanation is that, despite washing, streptomycin could accumulate in the host cytoplasm, where it gets in contact with bacteria and increase the selective pressure for streptomycin resistance. Although it is unlikely that the concentrations inside the washed host cells reached the minimum inhibitory concentration (MIC), sub-MIC levels of antibiotic have been shown to select for resistant mutants (Gullberg et al. 2011).

Each of the two RpsL mutations described above are almost in all cases accompanied by another mutation. RpsL43 is paired with a mutation in the ribosomal protein S4, RpsD, while RpsL88 is paired with a mutation in the co-chaperonin GroES. The specific mutation found in RpsD is a known compensatory mutation, which, in *E. coli*, restores an efficient translation to cells which also harbor the K43T mutation in RpsL (Björkman et al. 1999). The specific role of the co-chaperonin GroES (also referred to as HtpA in *Legionella*) in *Legionella* is not fully understood, but chaperonins in *Legionella* have been shown to play very different roles in the different phases of *Legionella* lifestyles. The chaperonin itself (GroEL/HtpB) appears to be exposed on the surface of the bacterial cell during infection and can trigger phagocytosis in non-phagocytosing cells, while it can redirect vesicular and organelle trafficking in the host cytoplasm when inside cells (Garduno et al. 2011). The function of the co-chaperonin GroES is less understood, but it is reasonable to assume that it co-translocates with GroEL. That specific mutation might also simply be compensatory, e.g. by helping to fold a potentially destabilized mutant RpsL protein.

RpsL, besides its function as a ribosomal protein, might trigger macrophage death, suggesting an adaptive role in host infection. Through pattern recognition receptors (PRRs), mammalian cells recognize bacterial invaders, and trigger both cellular apoptosis and inflammatory response (Akira et al. 2006). Many PRRs have known ligands (e.g. flagellin, needle of the type III secretion system) but some are still evasive. Experiments with mouse bone marrow-derived macrophages (BMDMs), Zhu et al. (2015) suggests that RpsL is responsible for macrophage death caused by *L. pneumophila*. Strikingly, in that study, only one of the two RpsL mutations found in our experiment (K88R) did not trigger cell death, allowing *Legionella* to replicate. Although we do not know how conserved this interaction is and whether human cells or amoebae possess a similar mechanism, it is interesting to note that 4 out of 6 of the Mac lineages display a fixed RpsL K88R mutation, while it is fixed in 2 Acas and 1 Alt lineages (other Acas lineages have two subpopulations with each one RpsL mutation). This suggests that, even in human macrophages, the K88R RpsL mutant might be less prone to trigger cell death. However, competition experiments do not seem to corroborate this observation: the RspL88/GroES genotype provides an important advantage in *A. castellanii* cells, but a slight disadvantage in U937 cells (**Figure 3**).

The RpsL43/RpsD mutation occurred twice in the lineages exposed to macrophages. We could not test the fitness of that genotype alone, since all the isolates we collected had one or more extra mutations. An isolate with an extra IolE mutation showed an increased fitness in *A. castellanii* cells but a decreased one in macrophages (**Figure 3**), a pattern similar to the RpsL88/GroES genotype. It is counterintuitive that a mutation with a fitness cost (in macrophages) would be fixed in all populations. A potential explanation is that the *Legionella* populations might have been exposed to streptomycin during the evolution experiment, setting a strong evolutionary pressure, but streptomycin (and thus the pressure) was not present at all during the competition experiment. These discordant results might be the result of the RpsL mutations being always beneficial in presence of (even traces of) streptomycin, and being beneficial in amoebae but detrimental in macrophages in the absence of streptomycin.

### An intergenic mutation in-between an ion-transport operon and a strain-specific gene

One single intergenic mutation was found to be fixed, upstream of two genes, lpp2016 and lpp2017 (**Figure 1**). The former gene, lpp2016, has no described function, and appears to be strain-specific in *L. pneumophila*, while the latter is involved in ion transport. From sequence analysis only it is not possible to determine whether this mutation would yield a selective advantage. However, it is possible that this mutation interferes with the regulation of one or both of the downstream genes. It is also possible that this intergenic space, unusually large (ca. 350 nt) might encode a small, non-coding RNA (ncRNA), which might have regulatory or even virulence functions. To the best of our knowledge, this specific intergenic region has not been shown to contain a ncRNA (Sahr et al. 2012). Recently, *L. pneumophila* has been shown to express and translocate sRNAs into its eukaryotic host, where it regulates the host innate immune response (Sahr et al. 2022).

### Recombination hotspots account for a large part of the mutations

*Legionella* genomes are known to experience extensive homologous recombination (David et al. 2017). Early studies estimated that the recombination (r) and mutation (m) rates should be similar (r/m = 0.9) (Coscollá and González-Candelas 2007; Vos and Didelot 2008). However, later estimates based on larger datasets suggest a much higher rate of recombination, with the vast majority of SNPs (96-99%, depending on the groups) arising from recombination (David et al. 2017). In our study, we identified five groups of homologous regions which harbor clusters of mutations. These regions are likely to represent hotspots of intra-genomic recombinations: eleven homing endonucleases, an ankyrin repeat-containing protein (lpp1100), one tetratricopeptide repeat (TPR) protein, a repeated protein containing a domain of unknown function (DUF1566), and three occurrences of a larger segment containing a transposase and a restriction endonuclease type II-like (DUF559). It should be noted that our experiment obviously excludes inter-genomic recombination, and that intra-genomic recombination among identical chromosomes yield no mutations. Only recombination between similar (but not identical) regions yields mutations, leading to an underestimation of the recombination rates.

The first group of recombination hotspots consists of homing endonucleases - found in all domains of life and in viruses - which are known to cause gene conversions between mobile genetic elements (Dunin-Horkawicz et al. 2006). This group of genes has been found to be enriched for in clinical sporadic strains (when compared to environmental strains) in *L. pneumophila* sequence type (ST) 1, suggesting that they contribute to the genetic diversity of ST1, the most prevalent ST (Mercante et al. 2018). The second group is an ankyrin repeat-containing protein (ANK). ANK motifs are typically eukaryotic motifs, and are found on many of the *Legionella* effectors (Habyarimana et al. 2008). This specific protein is not widespread in *Legionellales*, and is absent from most *L. pneumophila* strains, as well as from *L. longbeachae* (Gomez-Valero et al. 2011). It has been shown to be highly upregulated in transmission phase (Brüggemann et al. 2006; Jules and Buchrieser 2007). A distant homolog is however found in *L. drancourtii* LLAP12 (**Supplementary Figure S9**). TPR motifs, the third group of recombination hotspots, are known to be involved in protein-protein interactions in both eukaryotes and bacteria, and have been shown to be involved in virulence in *L. pneumophila* (Bandyopadhyay et al. 2012). The fourth hotspot is a non-repeated protein containing a domain of unknown function (DUF1566) which is widely present in Gammaproteobacteria. Finally, the fifth group of hotspots is constituted of three occurrences of a larger segment containing a transposase and a restriction endonuclease type II-like (DUF559). The first and fifth groups were recognized as repeated regions (David et al. 2017), and excluded from their recombination analysis. As for the other single-protein hotspots, they were not deemed repeated but were not detected as recombination hotspots.

About half (49.4%) of the mutations identified in our study occur in these five groups of mutational hotspots, but none of them got fixed. The majority of them occurred only at low frequency, and only mutations in three of the hotspot groups (lpp1100, GIY-YIG homing endonucleases, transposase/DUF559) reached 25% in frequency. However, the large number of intra-genomic recombination events that can be detected probably provides variation in the genome architecture and potentially contributes to the adaptability of *Legionella* genome.

The mutational patterns identified in this study follow the general principles of molecular evolution, for example displaying about twice as many transitions than transversions. Single-nucleotide polymorphisms are generally lowering the general G+C content of bacterial genomes (e.g. Hershberg and Petrov 2010; Ely 2021), while higher G+C content is generally believed to occur through GC-biased gene conversion (gBGC) (Lassalle et al. 2015). However, in our study, the AT to GC mutations unexpectedly accounted for almost half (49.4%) of the mutations altering the GC content, even after removing mutations occurring in recombination hotspots.

The pLPP plasmid found in *L. pneumophila* strain Paris (Cazalet et al. 2004) is sparsely found in ST1-strains (Durieux et al. 2019). Strikingly, although the plasmid is not essential, it has been retained in all of our lineages. This may be due to the presence of a toxin/antitoxin (TA) system on the pLPP plasmid: at least one PHD-RelE type TA system on the pLPP plasmid, according to the TADB (Shao et al. 2011) and TAsmania (Akarsu et al. 2019).

### Fitness of evolved strains

A puzzling observation in the fitness measurements performed is the discordance between our finding in the serial passage experiments of parallel and fixed mutations, which indicates a strong selective advantage, versus the lack of an advantage of the evolved clones (observed CI < 1) in competition experiments with the ancestral strains. At present we do not understand the underlying reason(s) for this discordance, but it should be noted that measuring fitness is a complex matter and in an experiment like the one described here, selection is acting at different levels. Compared to experiments in pure culture like the LTEE (Lenski 2017) where the maximum growth rate in exponential phase is paramount, other aspects of the life cycle of *Legionella* play a role. The time necessary to find a host, the ability to trigger endocytosis, the ability to resist digestion and the efficiency to recruit nutrients to the *Legionella*-containing vacuoles, the time required to exit lag phase and enter exponential phase, the efficiency to exit the host cells, are important in how many of each genotype make it to the next passage. Thus, the CI method used here is probably too crude to capture all these aspects, which might explain the sometimes-discordant results presented above. It is also possible that the fitness changes are non-transitive, and that competition experiments of the evolved strains versus the ancestral strain are not properly capturing such dynamics. Additional measures of fitness might help better understand the phenotypic effects of the mutations.

### Comparisons with other studies

Other studies have identified mutations specifically occurring within-host. In the first and - to our knowledge - only published evolution experiment with *L. pneumophila*, Ensminger *et al*. (2012) passage *L. pneumophila* in mouse macrophages for a year. This study identified, among others, mutations in the lysine synthesis pathway and in the flagellar regulation gene *fleN*. Surprisingly, there was no overlap between the mutations found in this study and ours. This might however also reflect the differences between the hosts and *Legionella* strains used in the two studies: while we used human cell lines and *A. castellanii* as hosts and *L. pneumophila* Paris, the other study cycled *L. pneumophila* LP01, derived from the Philadelphia strain, cycled in mouse-derived macrophages. A comparison with another study identifying mutation occurring in-patient (Leenheer et al. 2023) yielded one single match: in this study, the gene *lidB* (lpp2223/lpg2269) was found to be mutated twice independently in different outbreaks, while this gene is mutated in 13% of the population Mac_A at t2. The protein is an ATP-dependent helicase/deoxyribonuclease, possibly involved in replication. However, the evidence in both studies is too sheer to conclude a possible selective advantage of a mutated version of this protein without further functional studies.

### Early fixed mutations

An unexpected finding was the presence of multiple, apparently fixed mutations at the first time point (i.e. after 10-15 passages), some of which subsequently disappeared (**Figure 1**). We first discuss the probabilities of the same mutations appearing independently in various lineages, and then the fact that some of them disappeared, despite being apparently fixed.

The analysis of mutation rates and population sizes shows that parallel evolution in our setup is not unlikely. The mutation rates to streptomycin (mutations per cell per generation) measured by fluctuation assays on our ancestral strains (*L. pneumophila* Paris and the two fluorescently labeled strains) were in the range of 6.5 × 10^-10^. In comparison, mutation rates for bacteria range from 6 × 10^-11^ to 3 × 10^-8^ (Lynch et al. 2023), putting our isolates in the middle of the range. We calculated the probability of the appearance of a mutation providing resistance to streptomycin, given the number of generations and the mutation rates above, assuming no selection. Considering that streptomycin resistance is provided in most cases by two single RpsL mutations (K88R and K43T), the probability of gaining any given mutation is close to the probability of gaining a streptomycin resistance mutation. For an amoeba-exposed or a human cell-exposed population, the probability was 4% and 39% at the first time point (t1), while it reached 23% and 76% at the last time point (t3), respectively. This shows that it is not unlikely that single, identical mutations would appear parallelly in different populations. Given a strong enough selection, these mutations could have been quickly fixed in the population.

However, although other recurring mutations (e.g. in LerC or in lpp0833) are very likely to have occurred independently in different lines, the two paired mutations (Rpsl43/RpsD and Rpsl88/GroES), which always occurred together and in the same genetic background (SYFP2 and dTomato, respectively), have a higher likelihood to be the result of a spread of these two genotypes very early in the experiment. We had established very rigorous procedures to minimize the risks of cross contaminations: bacteria were grown in separate flasks and manipulated individually, always in a laminar flow cabinet, using filter tips when pipetting, and applying general sterility precautions when manipulating the strains. These two genotypes have rapidly taken over most of the populations, emphasizing their probable high selective value. Although a possible early contamination across lineages might have caused the spread of the RpsL mutants, we believe that the latter would be the only ones to be spread that way, and thus that the conclusions of the paper would not be affected.

To explain the fact that a few apparently fixed mutations disappeared at a later time point, we consider three possible explanations: contaminations across lineages, reversions, or not deep-enough sequencing. As mentioned above, we have established very rigorous procedures to minimize the risks of cross contaminations and thus favor the latter two hypotheses, either that the mutations reverted or that the sequencing missed the presence of the original allele. The apparently high selective value of some of the later mutations might explain these reversions or re-emergence of low-frequency wild-type alleles. An analysis of the reads mapping to the genes encoding the fluorescent markers (SYFP2 and dTomato) shows that in the populations harboring mutations considered fixed, and which therefore should contain only the reads mapping to only one of the fluorescent genes, a few reads of the other gene can be found (**Supplementary Figure S10**, see e.g. Acas_D or Alt_E). This suggests that the population still contains at least low frequencies of both populations, which might explain how apparently fixed mutations might disappear, should a more favorable mutation appear on the genome of the low-frequency genotype. It should be noted, however, that this later hypothesis is not particularly probable, as these favorable mutations are more likely to occur on the high-frequency genomes.

## Conclusion

In summary, the exact role, fate and histories of the mutations that occurred in the 18 lineages of *L. pneumophila* investigated here remains to be established. However, for both hosts, we identified potential adaptive, host-specific mutations. Lineages exposed to the amoebal host *A. castellanii* harbored mutations in the host-specific part of the LPS synthesis cluster. Lineages exposed to the human macrophage-derived cells displayed mutations in the LerC protein, a protein regulating the expression of protein effectors, potentially highlighting the different temporal regulation of protein effector secretion in the human host compared to *L. pneumophila*’s natural hosts. These results also show that experimental evolution can reveal bacterial genes specifically involved in the adaptation to certain hosts.

## Material and Methods

### Bacterial strains, host cell culture and media

*Legionella pneumophila* Paris was cultured in charcoal yeast extract (CYE) (1% ACES, 1% yeast extract, 0.2% charcoal, 1.5% agar, 0.025% Iron (III) pyrophosphate, 0.04% L-cysteine, pH 6.9) plates or ACES yeast extract (AYE) (1% ACES, 1% yeast extract, 0.025% Iron (III) pyrophosphate, 0.04% L-cysteine, pH 6.9) broth at 37℃, unless otherwise stated. When necessary, *Legionella* GVPC selective supplement (Oxoid) or 0.5 mM IPTG were also added to the media. The *L. pneumophila* strains used were tagged with a SYFP2 or dTomato fluorescence gene, under an IPTG-inducible promoter. A *L. pneumophila* Paris mutant harboring a deletion in the *dotA* gene was used as a negative control in infection experiments, as this strain is not able to replicate in host cells. The Δ*dotA* mutant was kindly gifted to us by Elizabeth Kay.

*Acanthamoeba castellanii* strain Neff (ATCC 30010) was cultured in Peptone Yeast Glucose (PYG) medium (ATCC 712 medium) in culture flasks at 30℃. For infections, *A. castellanii* was harvested, pelleted, and resuspended in LoFlo medium (Formedium, Norfolk, UK), which does not support *Legionella* growth.

Human monocyte-like U937 cells (ATCC-CRL-3253) were maintained in RPMI1640+GlutaMAX™ (Gibco) supplemented with 10% heat-inactivated fetal bovine serum (FBS) (Gibco) and 1% Penicillin-Streptomycin (Pen-Strep; Gibco), in a 37℃ incubator with 5% CO_2_. To induce differentiation into macrophage-like cells, the U937 culture was centrifuged at 200 × *g* for 5 min, resuspended in growth medium containing 50 ng ml^-1^ of phorbol 12-myristate 13-acetate (PMA) and incubated for 48h. The medium was replaced and cells were incubated for a further 48h. Before infection, cells were resuspended in RPMI 1640 without phenol red (Gibco) supplemented with 10% heat inactivated FBS and 1% GlutaMAX™ (Gibco). This medium also does not support *Legionella* growth.

### Passaging

*L. pneumophila* SYFP2 and dTomato patched on CYE for 48h, were resuspended in dH_2_O, the OD_600_ was measured and the cultures were diluted, before being mixed in equal proportion. Host cells were prepared as above, 2 × 10^6^ cells of *A. castellanii* or U937 were seeded in six T25 flasks, and challenged at an MOI of 0.25 (*A. castellanii*) and 2.5 (U937 macrophages). After inoculation, flasks were incubated at 30°C, or 37℃ for 3 days. At the end of the infection period, flasks were vortexed, and an aliquot of the culture was collected. These aliquots were centrifuged for 5 min at 200 × g to pellet the host cells. The endpoint concentration of *L. pneumophila* for each infection was estimated by measuring the OD_600_, and calculating the CFU based on a standard curve (OD_600_ vs CFU/ml) made for this purpose. The volume of bacteria needed to infect the hosts at the same MOIs was used to inoculate fresh host cultures. Thus, each new passage of *A. castellanii* was inoculated with 5 × 10⁵ CFUs, whereas the macrophages lineages were inoculated with 5 × 10⁶ CFUs. To confirm there was no contamination of the cultures, 10 µL of the cultures were plated on LB agar and CYE plates. *L. pneumophila* was passaged in this manner every 3-4 days, and at every 5th passage glycerol stocks were made by collecting all the infection supernatant (after pelleting the hosts) and centrifuging at 7000 × g for 15 minutes to pellet the bacteria. The resulting pellet was resuspended in AYE with 50% glycerol, and kept at −80℃.

### Measure of growth rates

To isolate individual clones, frozen stocks of the selected populations were streaked on CYE+GVPC plates and incubated for 72h. The bacterial lawn was scraped, resuspended in ultrapure water, and serial dilutions were made and plated on CYE+IPTG plates to obtain single colonies. Two colonies of each population were re-streaked, and glycerol stocks were made.

Fitness assays of the ancestor and evolved strains were performed both in AYE broth and in hosts. For growth in broth, a 24h patch of each colony was resuspended in AYE, the density (OD_600_) was measured and adjusted to 0.1 (ca. 2 × 10^8^ CFU/ml) with AYE+IPTG. Each culture was aliquoted in triplicate into a black 96 well plate, the growth rate was tracked by measuring the OD_600_, and fluorescence of SYFP2 (excitation: 508 nm, emission: 555 nm) or dTomato (excitation: 554 nm, emission: 635 nm) every 30 minutes for 24h hours using a Tecan Spark™ equipped with a monochromator. For in-host growth assays, bacteria and hosts were prepared as before, but here the 1 × 10^5^ host cells were seeded per well in a black 96-well plate and challenged at an MOI of 50. The infection media was supplemented with 0.5 mM IPTG. Intracellular replication was tracked by measuring the fluorescence every 30 mins, for 72h, as described above.

### Competition assays

Competition assays were performed in human monocyte-like cells (U937) and *A. castellanii* amoebae using a similar protocol to the one used by Ensminger et *al.* (2012). For in-amoeba competitions, evolved strains to be assayed were streaked onto CYE plates with IPTG (1 mM) and incubated at 37°C for three days. Biomass was taken from the lawn created by each strain and resuspended into either LoFlo medium (Formedium, Norfolk, UK) with 1mM IPTG (for infections in amoebae) or in RPMI without phenol red and with 1mM of IPTG (for infections in U937 cells). These solutions were diluted to an OD_600_ of 0.50 (around 5 × 10⁸ CFU/ml) with a Tecan Spark™ 10M plate reader. Each tested strain was mixed in equal parts with the ancestral strain (*L. pneumophila* Paris) with the other fluorescent tag: dTomato-tagged ancestral strain with SYFP2-tagged evolved strains, and SYFP2-tagged ancestral strains with dTomato-tagged evolved strain. The solutions were serially diluted down to 1/10th and a volume, depending on host species, was transferred to corresponding wells of 24-well plates containing either *A. castellanii* or U937 cells.

For competitions in amoebae, 200 µl of the mix of bacteria were incubated with 5 × 10⁵ *A. castellanii* cells in 2 ml of LoFlo medium with 1 mM of IPTG, to reach a multiplicity of infection (MOI) of 10. This MOI was chosen as it exceeds the minimal level of fluorescence detectable by the plate reader at the beginning of the assay. The plates were incubated at 30°C.

For competitions in macrophages, the U937 differentiated macrophages-like cells were seeded in the wells at 2.5 × 10⁵ cells per well, adding 100 µl of mixed evolved/ancestral bacterial strain solution to reach a MOI of 10. RPMI without phenol red but with 1mM of IPTG was used as medium for the infection of the bacterial inoculum solutions. Finally, the 24-well plates were incubated at 37°C with 5% CO_2_.

For both competitions, the ancestral dTomato- and SYFP2-tagged *L. pneumophila* were also competed on each plate, as a reference. All competitions were done in 5 replicates, for each pair of strains tested. The following controls were also performed on each plate: growing amoebae or U937 cells only (blank control), ancestral SYFP2-tagged strain and amoeba (SYFP2 positive control), ancestral dTomato-tagged strain and amoeba (dTomato positive control), and Δ*dotA* mutant (infection negative control, infection-defective). The *ΔdotA* was used as infection-negative control, CFU/ml counts were measured at the beginning and at the end of the competition experiment.

The fluorescence was measured with the Tecan 10M spark spectrofluorometer at an excitation/emission wavelength of 508/555 nm and 554/635 nm to measure SYFP2 and dTomato fluorescence, respectively. Fluorescence was measured at 0 hours, 40 hours, and 64 hours post infection. These time points correspond to the beginning of the infection, to the beginning of the exponential phase, and the beginning of the post-exponential phase of growth, respectively, in our experimental setup. The time points were determined by measuring growth curves and optimizing the conditions (see **Supplementary Figure S7**). Fluorescence data was used to calculate a competition index (Ensminger et al. 2012). The fluorescence measurements were blanked with the fluorescence values of the amoeba-LoFlo or U937-RPMI control wells. Competition index (CI) is calculated as the ratio of the fluorescence of the mutant (evolved) strain to the fluorescence of the wild type (ancestor) strain at t1, divided by the same ratio at t0:

CI = [F(mut)_t1_/F(WT)_t1_]/[F(mut)_t0_/F(WT)_t0_] Where:

F(mut)_t1_ = Fluorescence of mutant at end time point (t1)

F(WT)_t1_ = Fluorescence of wildtype at end time point (t1)

F(mut)_t0_ = Fluorescence of mutant at beginning time point (t0)

F(WT)_t0_ = Fluorescence of wildtype at beginning time point (t0)

We normalized the CI data of each strain tested by dividing their CIs by the average CI of the SYFP2-tagged ancestor against the dTomato-tagged ancestor. This allowed us to compare CIs across plates. The competition index reflects the ratio of the number of mutant cells to the number of ancestral cells, produced between t0 and t1.

### Fluctuation assays to determine mutation rates

Fluctuation assays were carried out according to the recommendations of Lang (2018) and Rosche and Foster (2000). Briefly, the tested strains (*L. pneumophila* Paris WT and the two SYFP2 and dTomato-tagged mutants) were inoculated in AYE broth and diluted to reach an OD_600_ of 0.50, corresponding to ca. 5 × 10^8^ CFU/ml. The solutions were diluted 1/1000, by adding 5 µl of this solution to 5 ml of fresh AYE, and vortexed. From these, 100 µl were used to inoculate 2 ml of AYE into 50-ml Falcon tubes. This was repeated 72 times for each strain tested. The tubes were incubated at 37°C for three days. After growth, 150 µl of the solution was used to count CFU/ml numbers using the serial-dilution spot plate method (Wang et al. 2017). The rest of the solution was centrifuged at 6000 × *g* and most of the supernatant removed. The bacterial pellet was resuspended in the remaining supernatant and plated on CYE containing 6 µg/ml streptomycin, which is around 8x the minimum inhibitory concentration (MIC). The plates were incubated for 3-4 days and total and resistant colonies were counted. Mutation rates were calculated using the web-based Fluctuation AnaLysis CalculatOR (FALCOR) (Hall et al. 2009). The MSS Maximum Likelihood Method was used to calculate mutation rates using the “Group all data” option.

### Identification of *rpsL* mutations in streptomycin-resistant strains

To survey the various mutations contributing to streptomycin resistance, we picked a total of 40 streptomycin-resistant colonies from the fluctuation assay and sequenced their *rpsL* gene, known to confer resistance mutations to streptomycin (Zhu et al. 2015). This was done by amplifying a 300-bp segment of the *rpsL* gene using the following primers: forward 5’-AAGAAAGCCTCGTGTGGACG-3 and reverse 5’-TCGGTCGTTCACTCCTGAAG-3’. The DreamTAq Green PCR Master mix (ThermoFisher Scientific) was used to amplify the gene and the manufacturer’s instructions were used for PCR reactions with an annealing temperature of 59.6°C and 25 cycles. PCR reactions were checked on 1% agarose gel, and then purified using the GeneJET DNA Cleanup Micro kit (ThermoScientific^TM^). The purified fragments were sent for Sanger sequencing to identify mutations.

### Streptomycin susceptibility testing

The level of streptomycin resistance of the ancestral strains and evolved isolates was evaluated using an E-test on CYE+IPTG agar plates. The minimum inhibitory concentration (MIC) was recorded as the lowest antibiotic concentration at which the zone of inhibition intersected the E-test strip; for many isolates there was a zone with haze of growth around the strip, here the MIC was read the same way, but the MIC at the start at the hazy zone was also recorded (**Supplementary Figure S5**).

The selected *Legionella* strains were grown on CYE+GVPC plates for 48-72h; a loop of bacteria was suspended in dH_2_O, and the density was adjusted to OD_600_= 0.05 (ca. 1 × 10^8^ CFU). Then 1 ml of the *Legionella* suspension was added to 5 ml of 0.5% (soft) agar solution, the mixture was gently vortexed, to avoid the formation of bubbles, and poured onto a CYE+IPTG agar plate. The plates were left to solidify for 10-15 min before applying the streptomycin E-test strip (AB-BIODISK, bioMérieux). The plates were then incubated at 37°C, and the results were taken at 48h.

### Population and clone sequencing

In order to identify the mutations that arose from the evolution experiment in the different lineages, we sequenced populations at passages 10 and 65 for the *A. castellanii* lineage, passages 10 and 49 for the alternating lineage, and passages 15 and 38 for the macrophage (U937) lineage (**Supplementary Table 1**). All replicates from each lineage were sequenced. All clones involved in competition experiments were also sequenced, to confirm that other mutations did not occur (**Supplementary Table 3**).

Briefly, the frozen stocks of the different lineages were plated on CYE agar plates and incubated at 37°C for 3 days. After incubation, the bacterial growth was scraped off from the plate and suspended in 5 ml of distilled water. The cultures were centrifuged at 6000 × *g* for 5 minutes and the supernatant was removed. Centrifugation and supernatant removal were repeated. The DNA was extracted using the DNeasy blood and tissue kit (Qiagen), using the pretreatment for Gram-negative bacteria, following the manufacturer’s instructions, with the following exceptions: the lysis incubation period was increased to three hours, 4 µl of 10 mg/ml of RNase A were used, and the DNA was eluted in water. Next, the DNA samples were prepared for barcoding and sequencing using the Nextera XT kit from Illumina. The manufacturer’s instructions were followed to prepare the libraries.

### Variant calling

Reads were processed for quality using fastp version 0.23.2 (Chen et al. 2018). Adapters were trimmed from the reads using the --detect_adapter_for_pe option and the --overrepresentation_analysis, --correction, and --cut_right options were used for quality trimming. Trimmed reads were deposited at ENA under study accession number PRJEB82630. Variant calling using the clean and processed reads was done using breseq version 0.36.1 (Deatherage and Barrick 2014). In brief, breseq calculates a Bayesian score that all disagreements with the consensus at each position derive from sequencing errors only. It also filters out potential mutations where there is a strand bias, a quality bias, and positions in homopolymeric stretches. Finally, it only calls variants with at least 5% frequency in the population. For population sequencing, the --polymorphisms-prediction option was used for variant calling.

One population where the hosts were alternated (Alt_D) displayed a large excess of mutations at passage 10 (t1): while the median number of mutations is 28.5 per population, Alt_D displayed 989 mutations. A transient contamination from another *L. pneumophila* strain could be identified in this population. The distribution of the mutation frequency showed a clear peak around 6% (**Supplementary Figure S11**). In this population only, all mutations with a frequency lower than 20% were filtered out, thus retaining only 15 mutations, of which none were fixed.

Mutations occurring in regions containing many tandem repeats were filtered out.

For coverage analysis of the reads mapping to the genes encoding fluorescent genes, the trimmed reads were mapped to the sequence of either gene using bowtie2 v2.4.4 (Langmead and Salzberg 2012), removing bad reads with --qc-filter.

### Statistical analysis and visualization

All statistics and most figures in this contribution were performed in R (R Development Core Team 2024), using the tidyverse (Wickham et al. 2019) and, in particular, the ggplot2 package (Wickham 2009: 2).

### Structure prediction of the LerC mutants

Alternative gene prediction of the *lerC* mutants (lerC_46 and lerC_61) was performed by inputting the mutant sequence of the gene in prodigal 2.6.3 (Hyatt et al. 2010), with default options. Structures were predicted using AlphaFold3 (Abramson et al. 2024) then superimposed using PyMol v2.5.4 (Schrödinger, LLC 2015).

## Data availability

Population sequencing data is available at ENA under Study accession number PRJEB82630.

## Acknowledgments

The authors would like to thank Chayan Kumar Saha, for helping with the LerC mutant structures.

This study was supported by grants from the Swedish Research Council [2017-03709 to L.G.], the Carl Trygger Foundation [CTS 15:184, CTS 17:178 to L.G.], the Fonds de recherche du Québec [FRQNT B3X Postdoctoral Scholarship to K.P.] and the Natural Sciences and Engineering Research Council of Canada (NSERC) [Postdoctoral Scholarship to K.P]. The funders had no role in the design of the study.

## Supplementary Material

### Supplementary Tables

**Supplementary Table 1:** Population sequencing and number of generations in the different hosts.

| Host | Abbrev. | t1<br>Passages | N generations | t2<br>Passages | N generations | t3<br>Passages | N generations |
| --- | --- | --- | --- | --- | --- | --- | --- |
| *A. castellanii* | Acas | p10 | 128 +/- 7 | p40 | 517 +/- 9 | p65 | 798 +/- 9 |
| Alternation | Alt | p10 | 124 +/- 5 | p49 | 481 +/- 13 | p85 | 799 +/- 13 |
| U937 cells | Mac | p15 | 113 +/- 5 | p38 | 270 +/- 10 | p65 | 444 +/- 40 |

**Supplementary Table 2:**
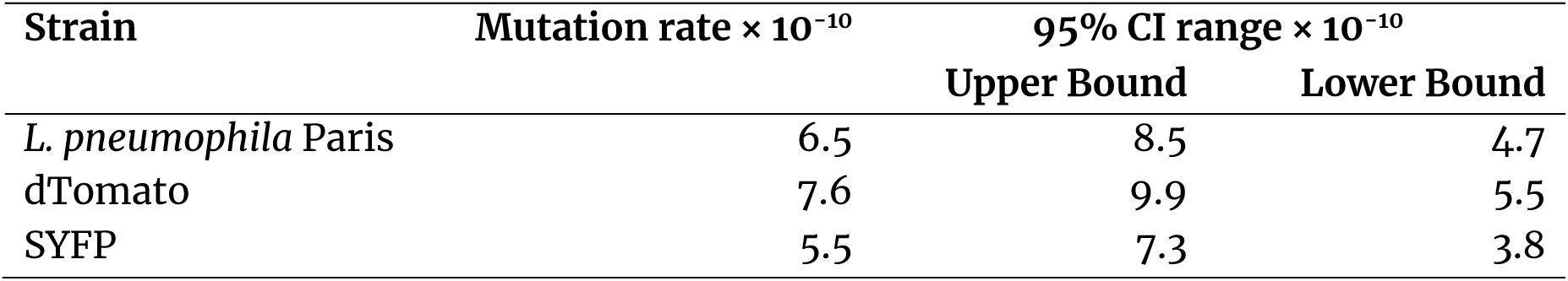
Mutation rates (mutations per cell per cell division or generation) measured for *L. pneumophila* Paris, dTomato-, and SYFP2-labeled strains. Mutation rates were measured using streptomycin as the selection antibiotic and then calculated using MSS Maximum Likelihood method. Mutations rates are expressed per cell per generation per billion. CI, Confidence Interval.

**Supplementary Table 3:**
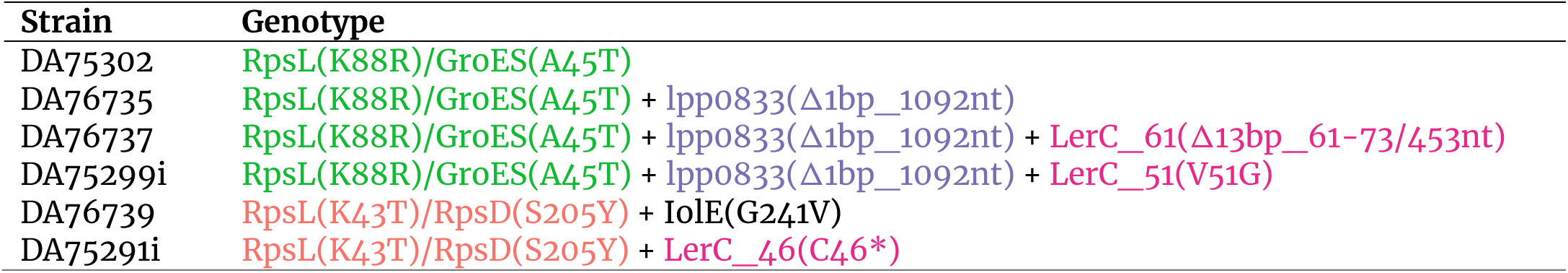
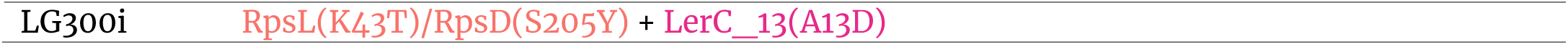
Clones and mutations involved in competition experiments. Genes and mutations are color-coded.

### Supplementary Figures

**Supplementary Figure 1:**
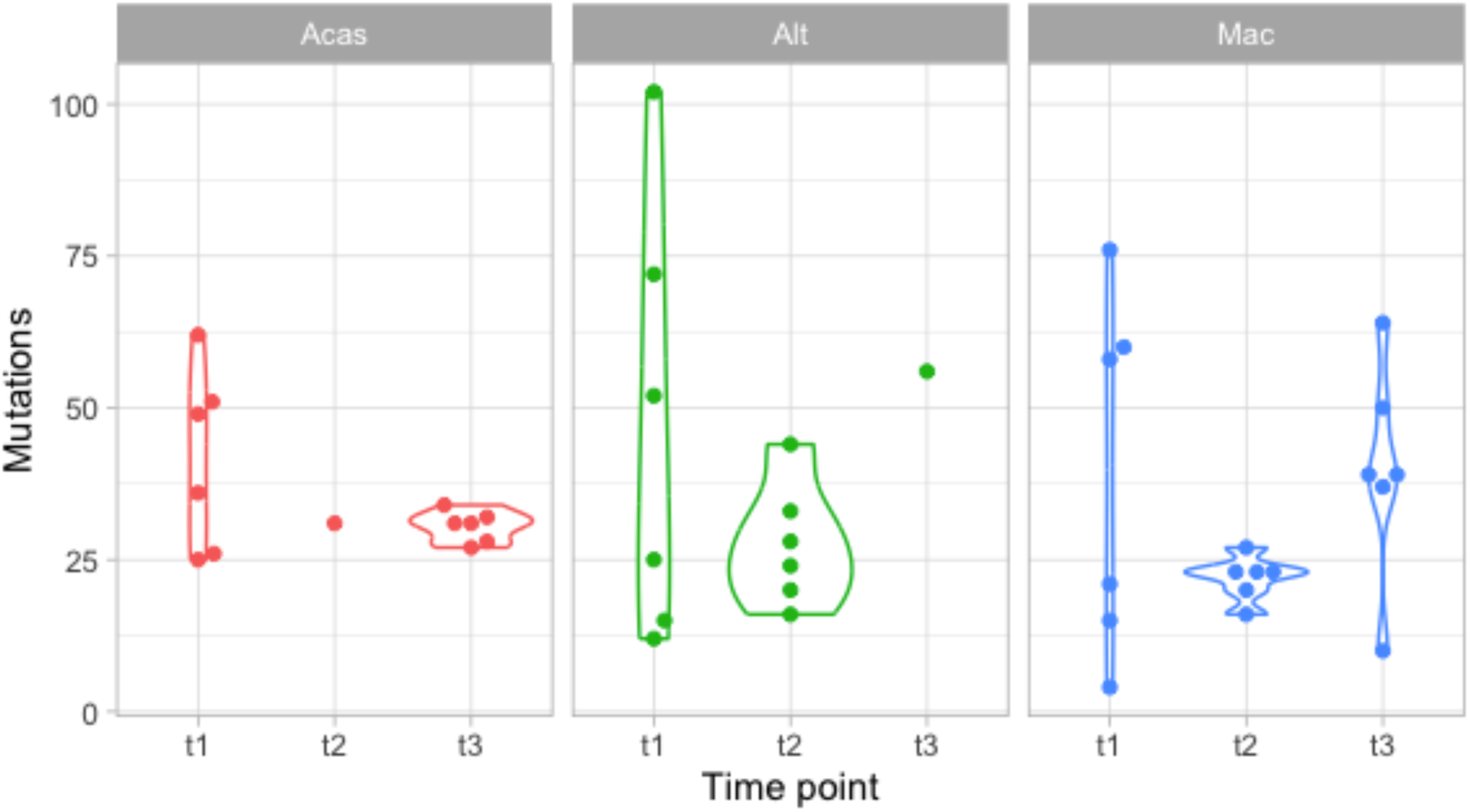
Distribution of mutations per host and time point. Each point is a lineage. Only one Acas population and was sequenced at t2, and only one Alt population was sequenced at t3.

**Supplementary Figure 2:**
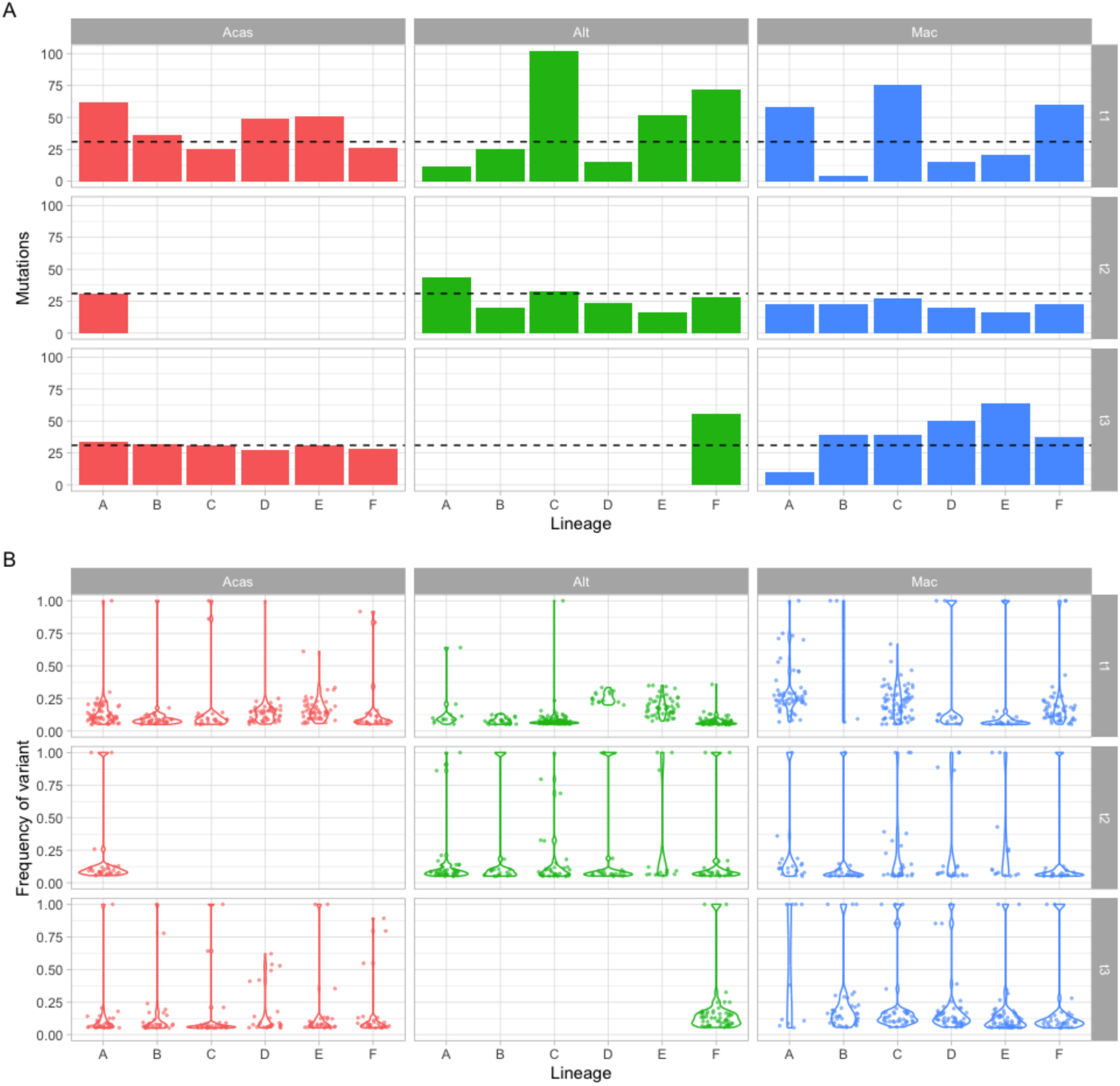
Distribution of mutation numbers and frequencies. **(A)**: Number of mutations with frequency > 5% per host (Acas, red; Alt, green; Mac, blue), lineage (A-F, x-axis) and time point (t1-t3). The median number of mutations per population (32.5) is shown with a dashed black line. **(B)**: Frequency (y-axis) of mutations per host, lineage and time point. Each dot represents a mutation, and its position on the y-axis represents its frequency. Violin plots are superimposed on the dots to provide a representation of the distribution.

**Supplementary Figure 3:**
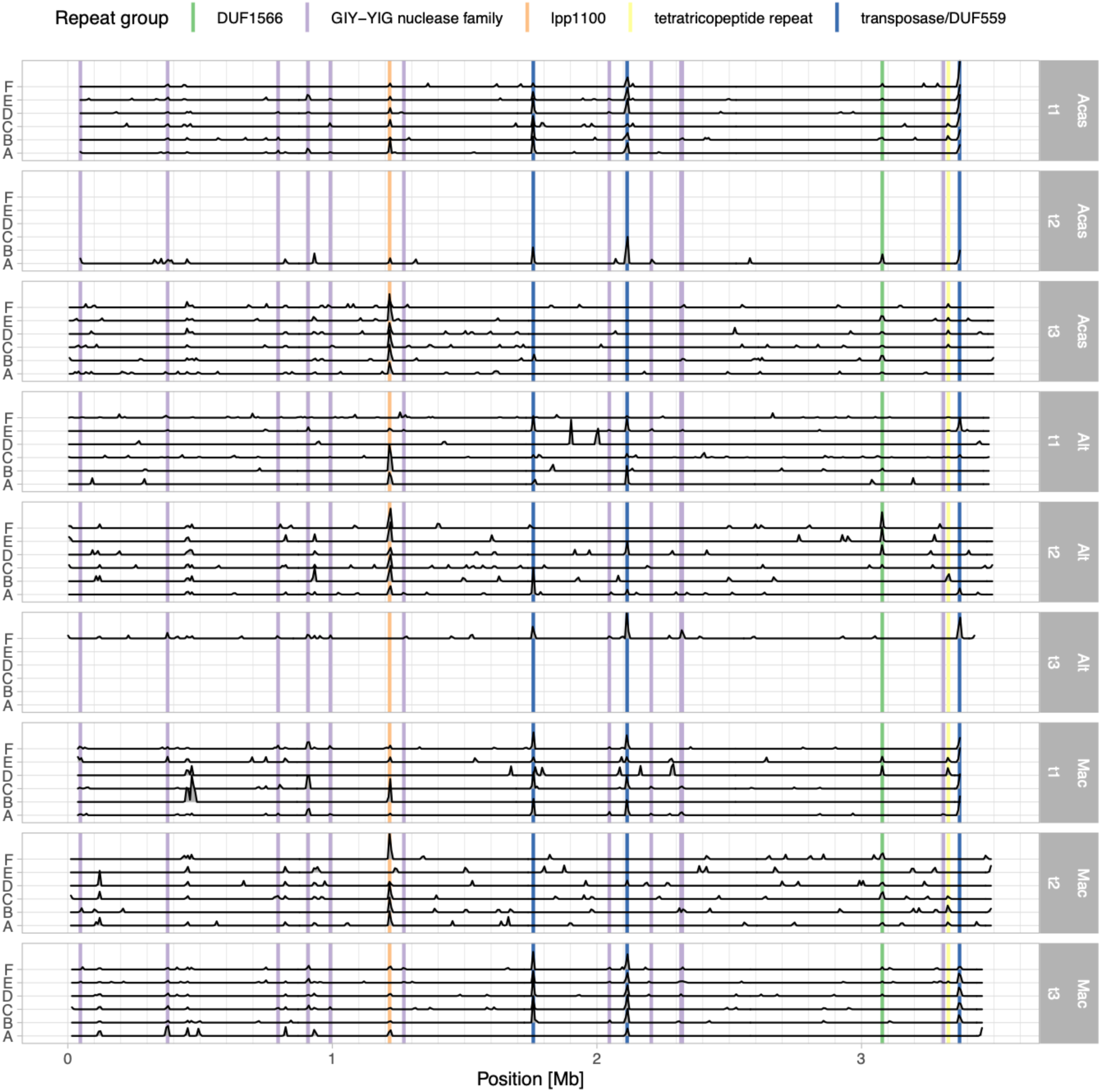
Distribution of mutations along the *L. pneumophila* genome. Each line corresponds to a separate population. The host and timepoint is indicated on the right, the lineage on the left. The position of genes containing repeats or being present in different parts of the genome are indicated with lines.

**Supplementary Figure 4:**
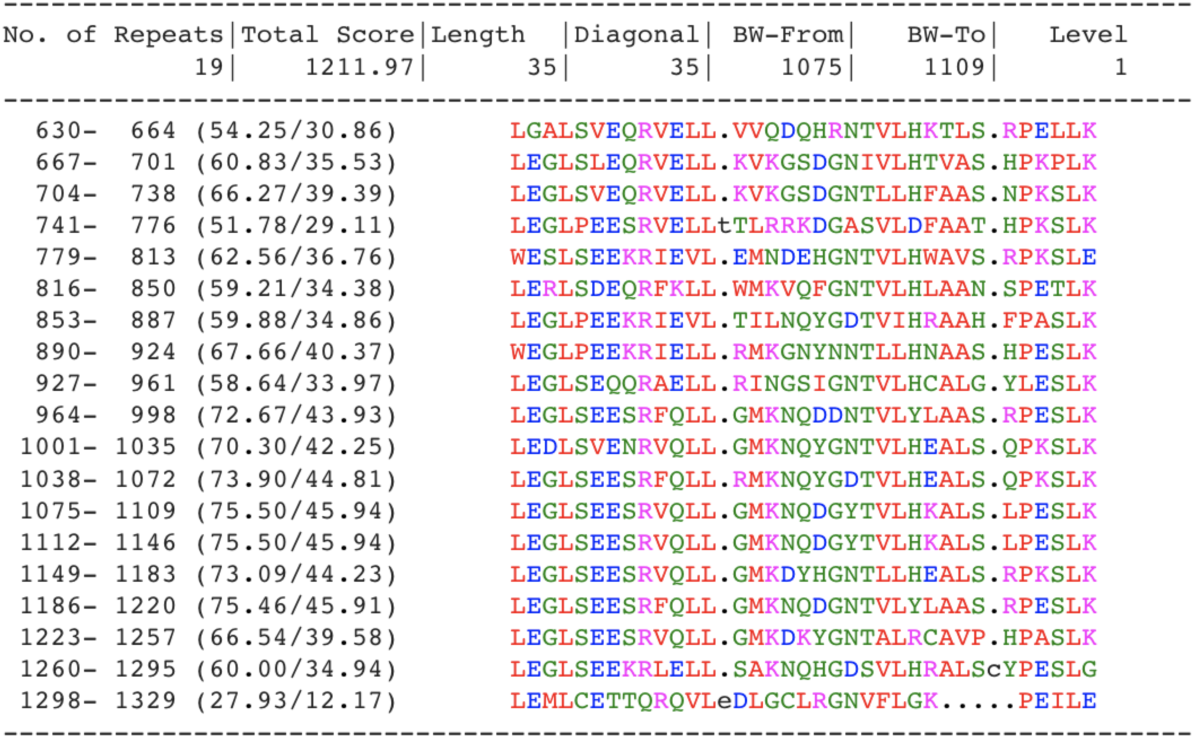
Repeats in ankyrin-repeat containing protein lpp1100. The analysis was performed using RADAR (Heger and Holm 2000).

**Supplementary Figure 5:**
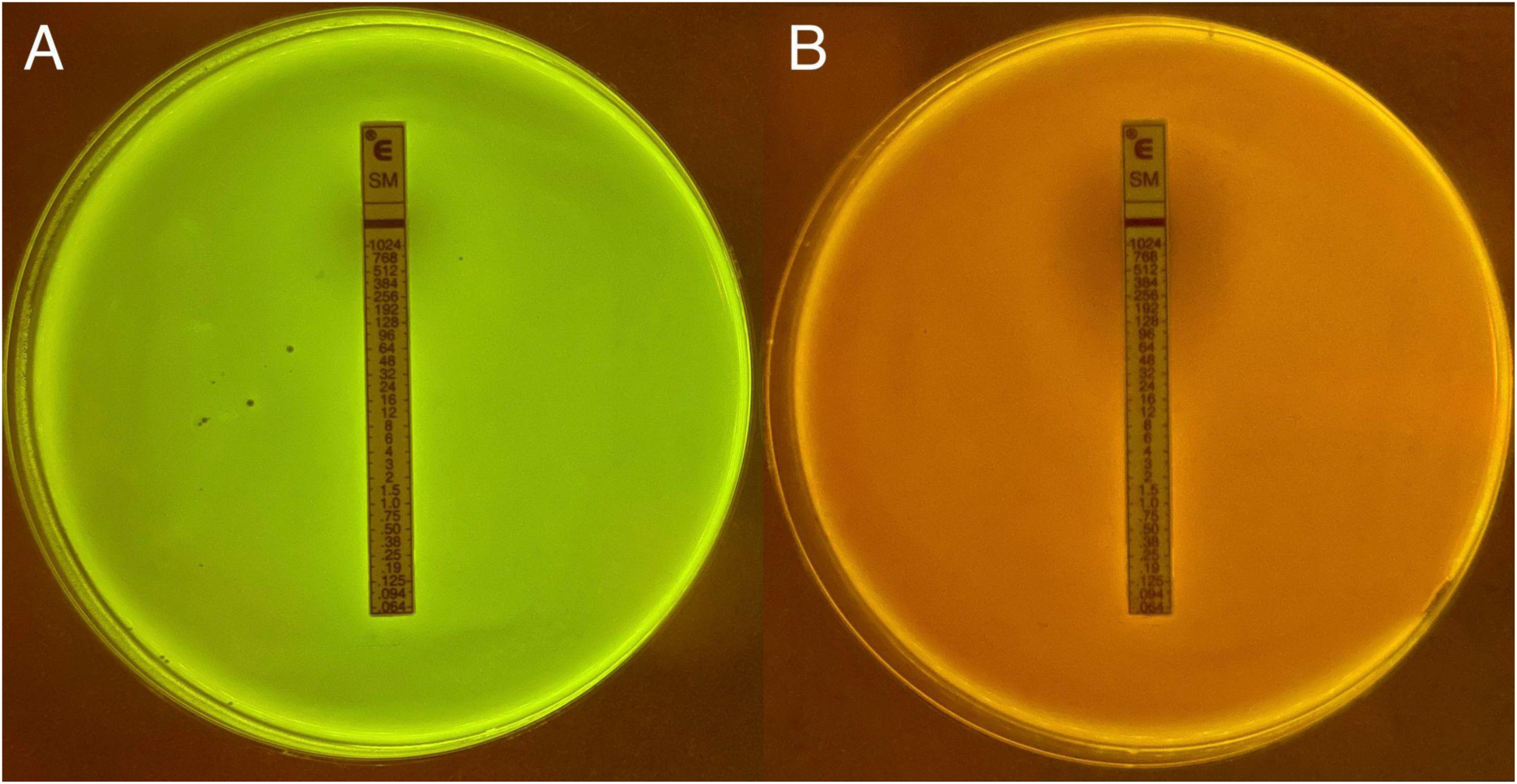
Resistance to streptomycin of two *Legionella* clones bearing different mutations in the *rpsL* gene. (A) *L. pneumophila* with the Rpsl43/RpsD genotype, in a SYFP2 background. (B) *L. pneumophila* with the Rpsl88/GroES genotype, in a dTomato background. Although the bacteria grow on the whole plate, including at the maximum streptomycin concentration (1024 μg/ml), a larger halo zone can be observed in B (to ~96 μg/ml) than in A (to ~768 μg/ml).

**Supplementary Figure 6:**
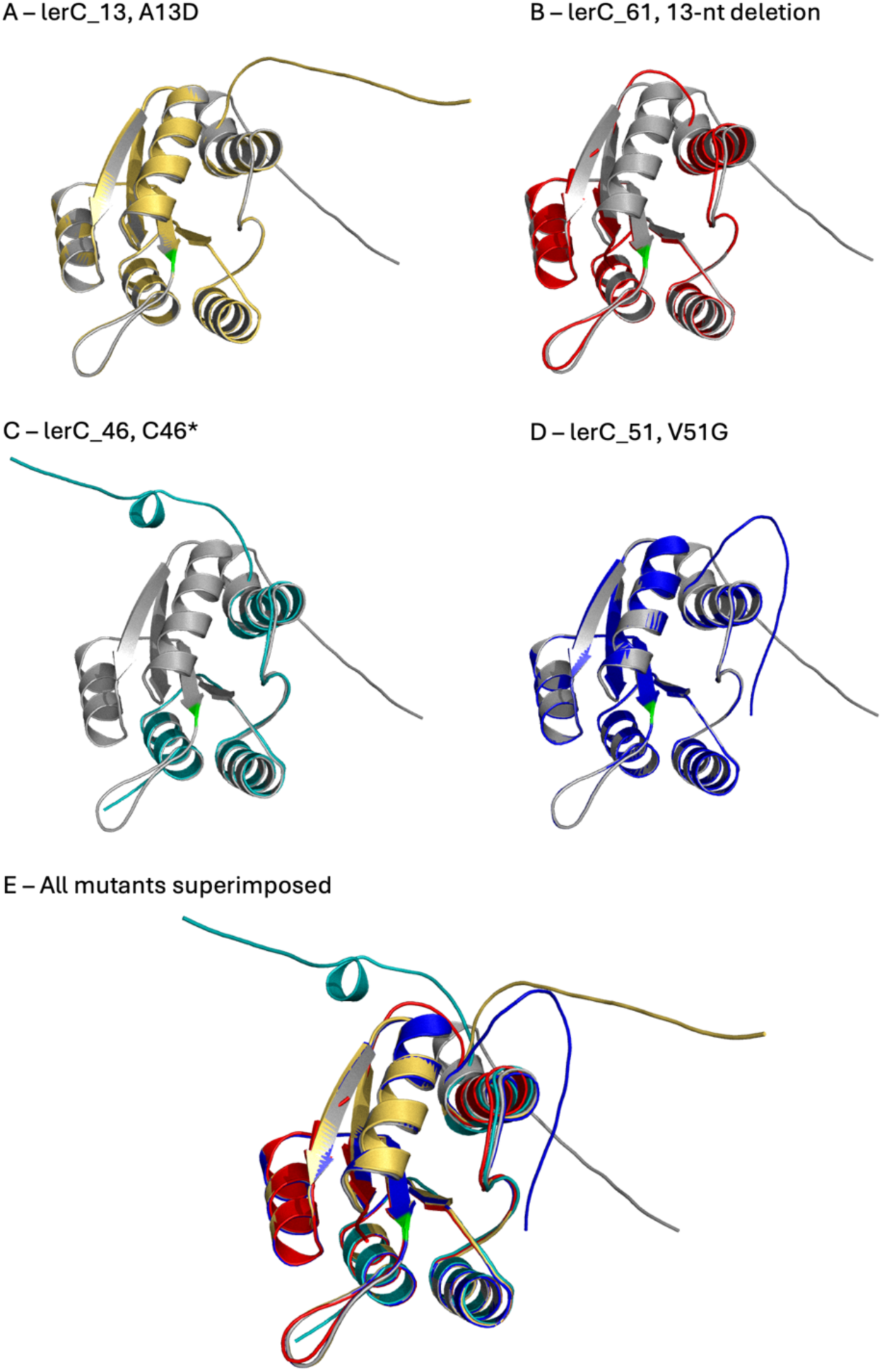
Structure of the *lerC* mutants, as predicted by AlphaFold3. The predicted wild-type LerC protein is shown in grey on all panels, lerC_13 in yellow (**A, E**), lerC_61 in red (**B, E**), lerC_46 in teal (**C, E**), and lerC_51 in blue (**D, E**). The location of the conserved aspartic acid (D53) is shown in green.

**Supplementary Figure 7:**
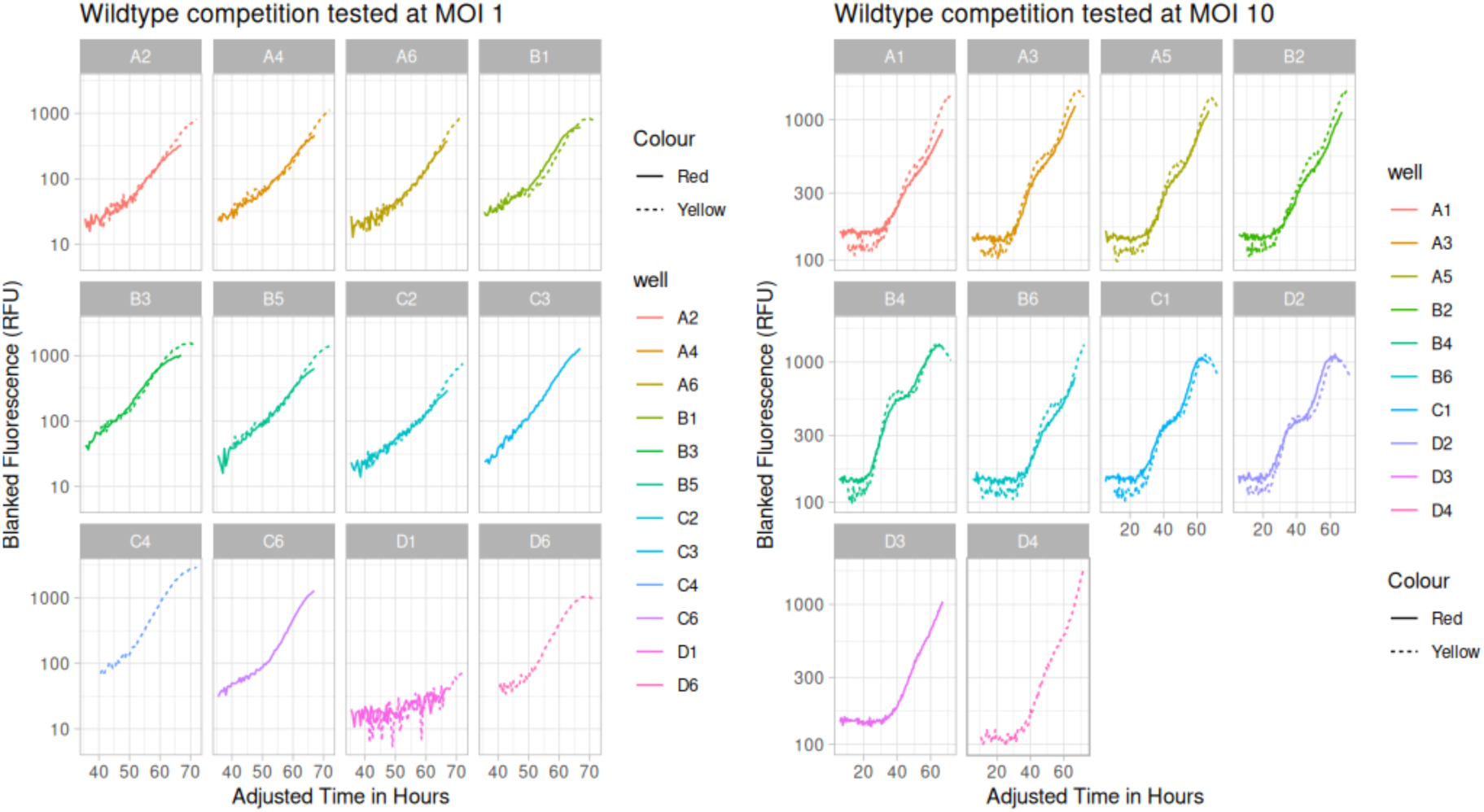
Competition experiments between selected isolates. Competitions of *L. pneumophila* strain Paris marked with dTomato (Red) and SYFP2 (Yellow) in an amoeba infection model at MOI 1 and MOI 10.

**Supplementary Figure 8:**
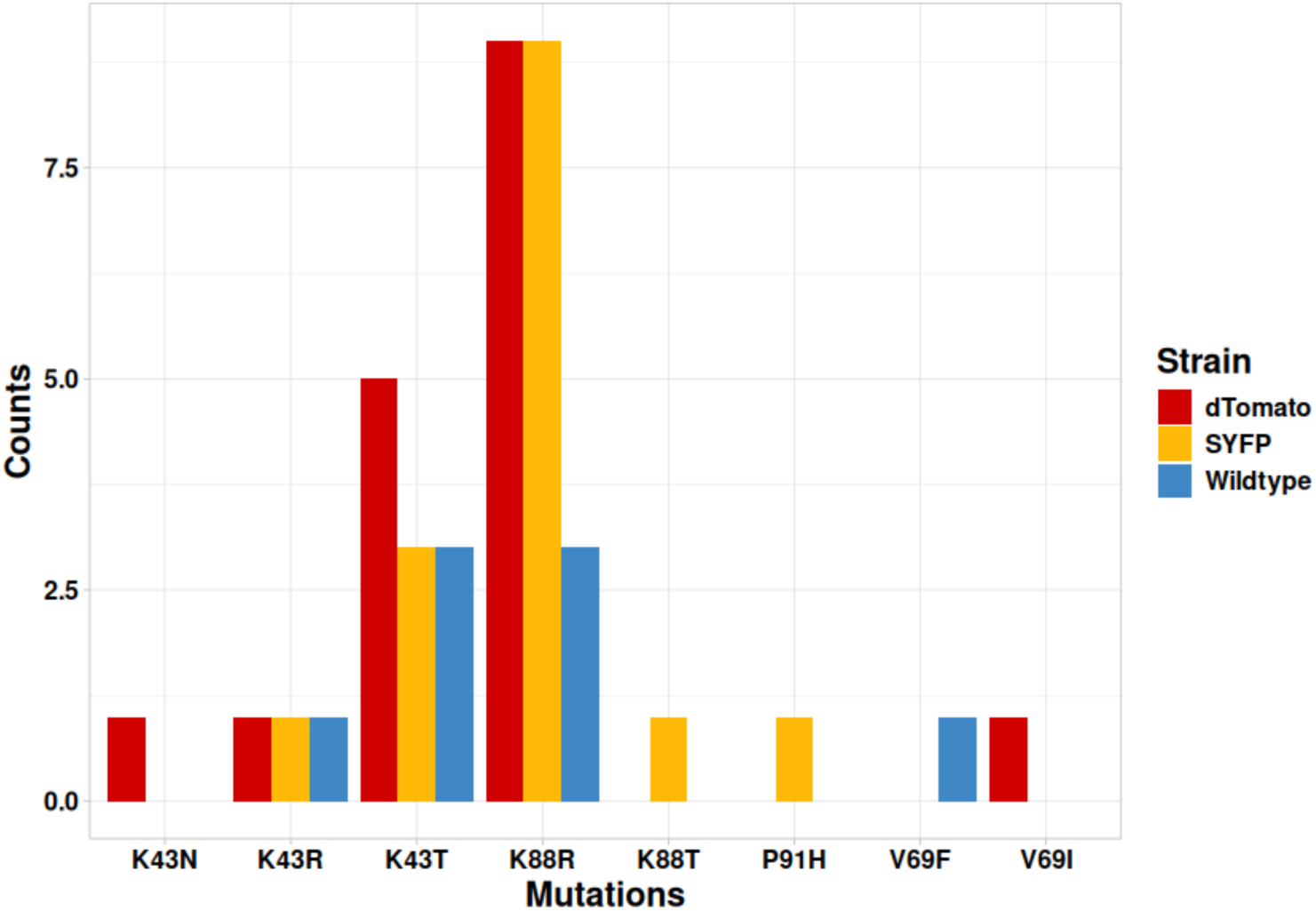
Distribution and diversity of non-synonymous mutations in the *rpsL* gene for the three ancestral strains (*L. pneumophila* Paris, SYFP, and dTomato). The x-axis represents the amino acid residues switched through the mutation and their position in the RpsL sequence.

**Supplementary Figure 9:**
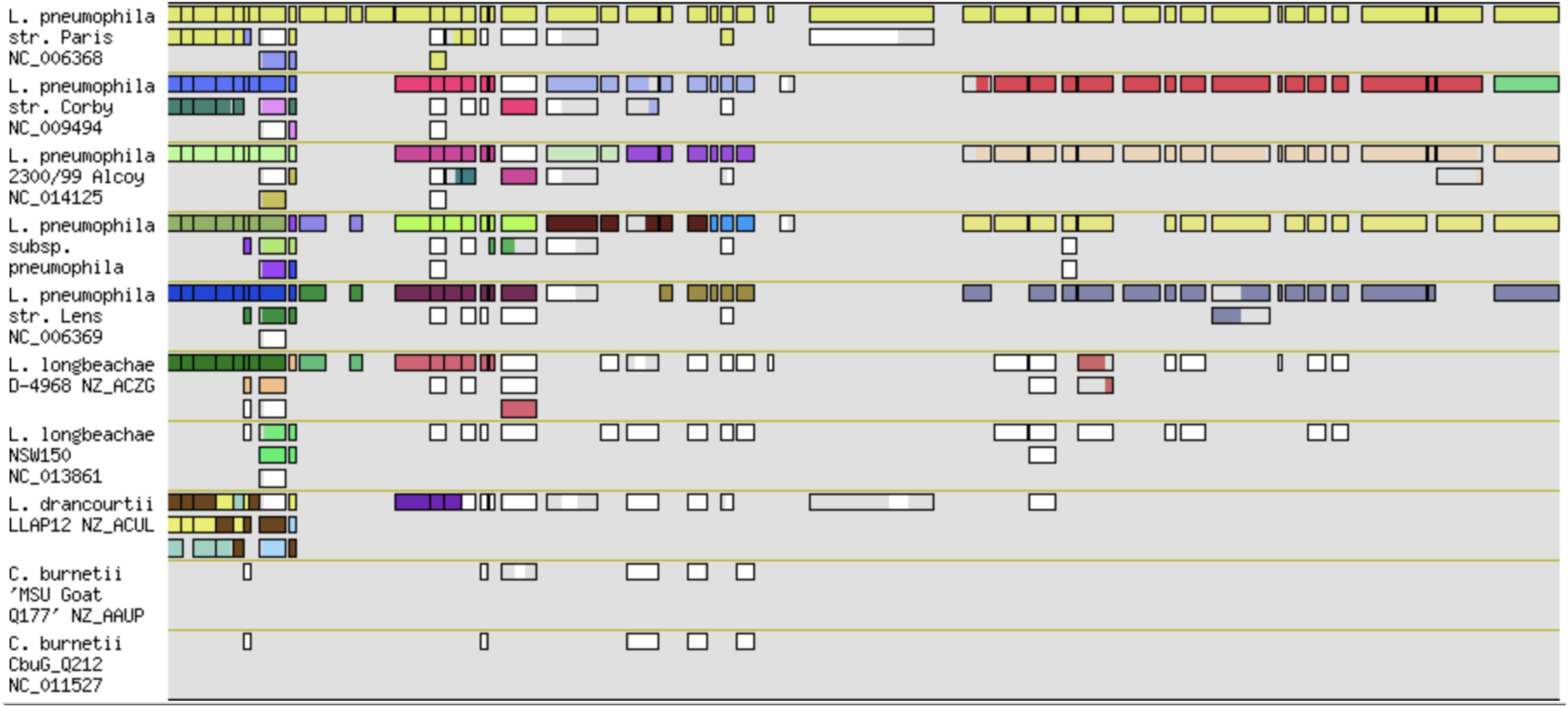
Synteny plot of *Legionella* and *Coxiella* species, centered around lpp1100 (LPP_RS05510) in *L. pneumophila* Paris (upper row). The other rows Colors indicate groups of genes colocalized in a specific genome. Figure obtained with MAGE (Vallenet et al. 2006).

**Supplementary Figure 10:**
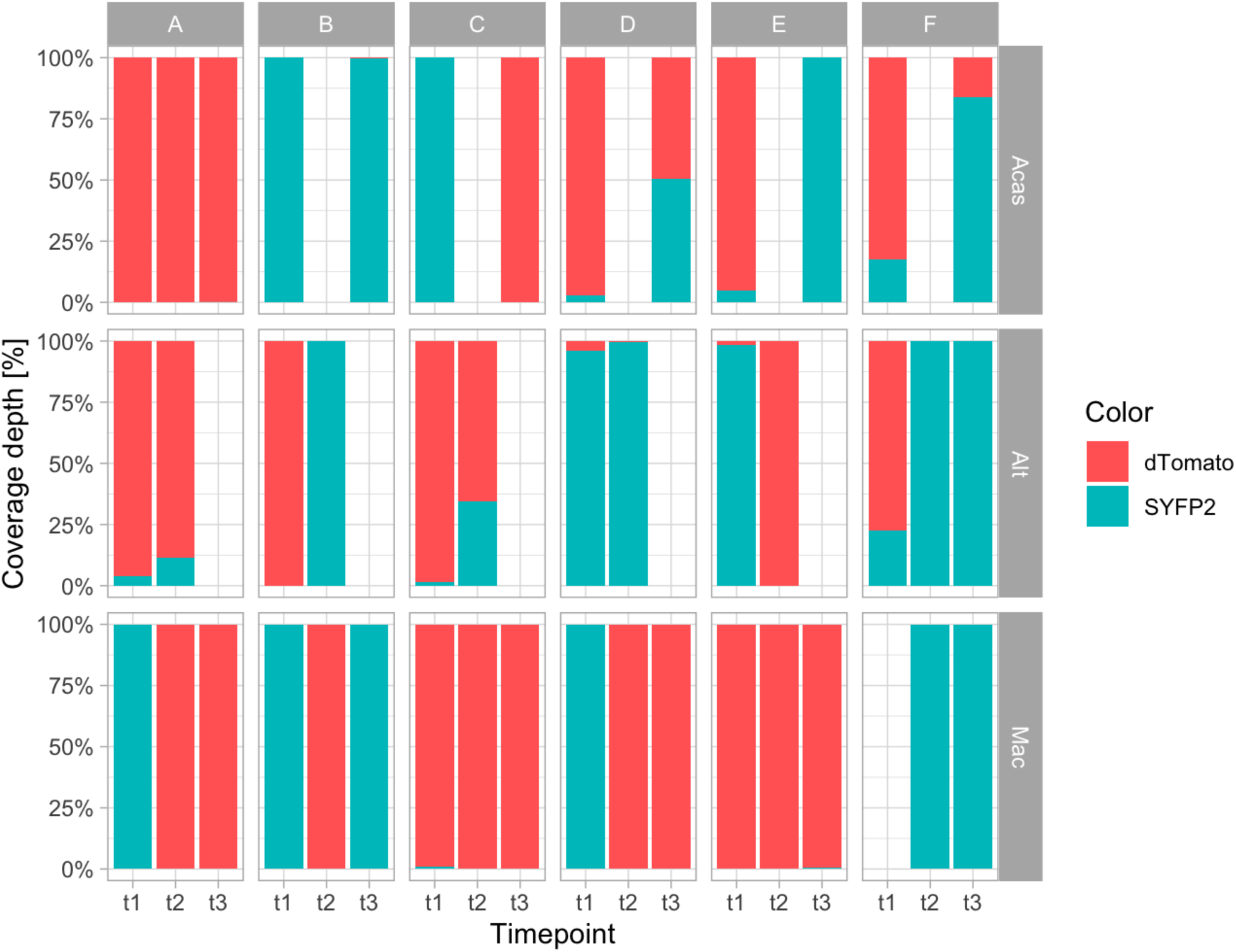
Proportion of SYFP2- and dTomato-marked *L. pneumophila* in each lineage at the 3 time points. At t0, each population consisted of an equal mix of each ancestor. The proportion at the other time points was estimated by mapping the reads obtained from population sequencing to the genes encoding the two fluorescent proteins (dTomato, red; SYFP2, green).

**Supplementary Figure 11:**
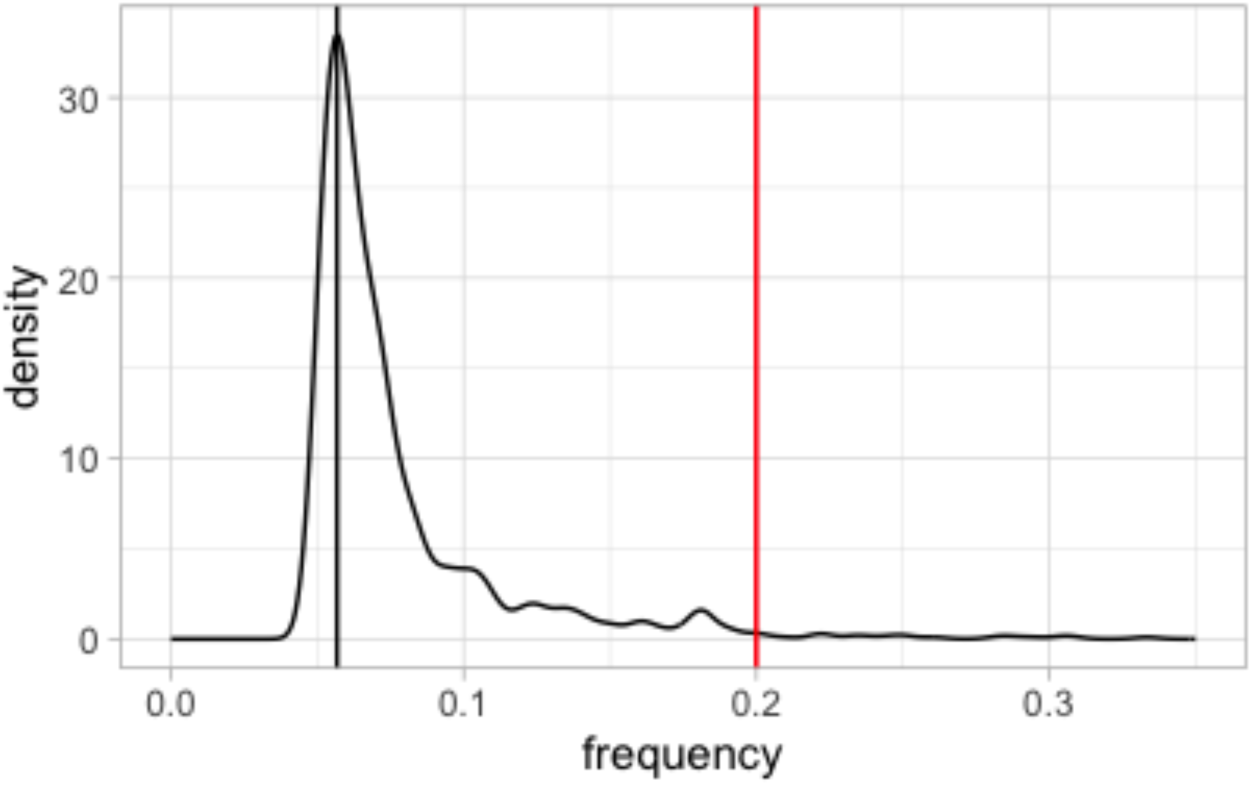
Distribution of mutation frequencies in the Alt D population, which counted 991 mutations. The black vertical line marks the maximum of the density curve, at 5.7%, while the red line marks the chosen 20% cut-off, under which mutations were discarded.

